# DNA binding contributes to dominant-negative effects of a catalytically inactive RecQ4-family helicase

**DOI:** 10.64898/2026.02.13.705743

**Authors:** Robert H. Simmons, Faith E. McDevitt, Alexandra Hurlock, Michael E. Kumcu, Matthew L. Bochman

## Abstract

DNA inter-strand crosslinks (ICLs) are highly cytotoxic lesions that require coordinated processing for repair. The RecQ4-family helicase Hrq1 promotes ICL repair in *Saccharomyces cerevisiae*, yet the mechanistic basis of its function remains unclear. Notably, the catalytically inactive *hrq1-K318A* allele confers greater sensitivity to ICL-inducing agents than deletion of *HRQ1*, suggesting a dominant-negative effect. To define the basis of this phenotype, we performed a genetic suppressor screen combined with biochemical and structural analyses. Spontaneous suppressors of *hrq1-K318A* sensitivity were overwhelmingly intragenic second-site mutations, many of which are predicted to destabilize the protein or impair its ability to bind DNA. In all cases, these mutations alleviated the dominant-negative repair defect. Biochemical characterization of representative mutants, including a rationally designed DNA-binding mutant, demonstrated that disruption of DNA binding suppresses *hrq1-K318A* toxicity even when protein stability is retained. These findings support a model in which DNA engagement by a catalytically inactive RecQ4-family helicase contributes to dominant-negative interference with DNA repair. More broadly, this work provides insight into how incomplete loss-of-function alleles of human *RECQL4* may disrupt genome maintenance pathways.

**GRAPHICAL ABSTRACT:** 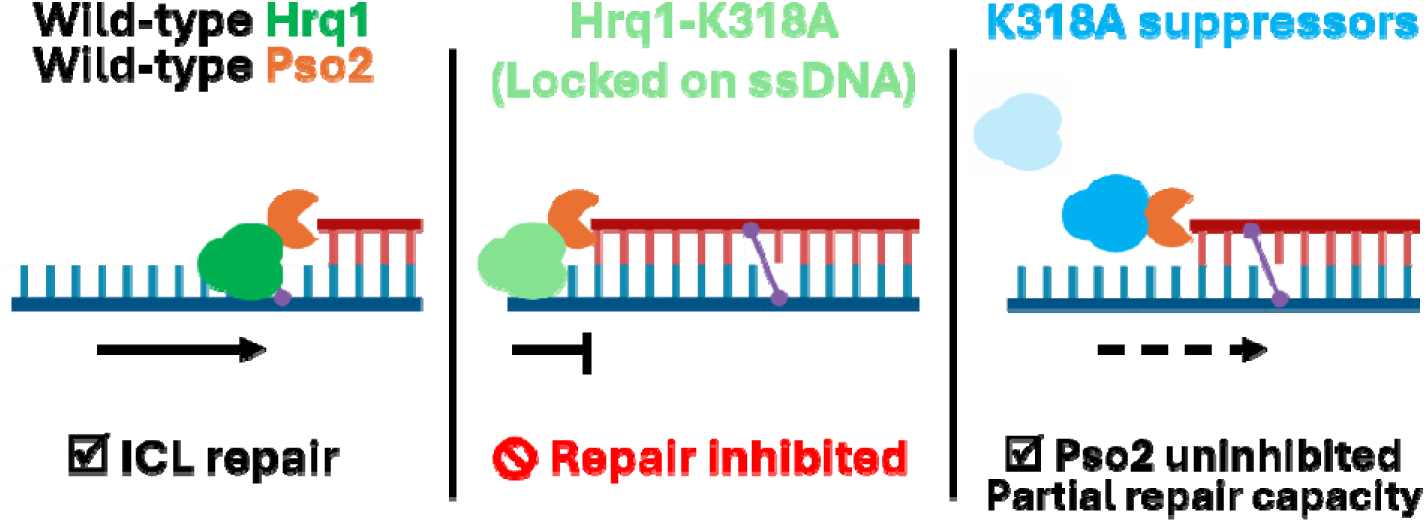

## INTRODUCTION

The RecQ family of helicases is an evolutionarily conserved group of enzymes that are found across all domains of life and have well-characterized and important roles in DNA replication, recombination, repair, transcription (1). The human genome encodes five RecQ helicases (RECQL1, WRN, BLM, RECQL4, and RECQL5), four of which cause autosomal recessive genetic diseases when mutated (2–5). These enzymes have also recently emergent targets for cancer therapeutics and diagnostics (1,6). For example, the WRN helicase has been identified as a synthetic lethal target for cancers that display microsatellite instability–high (MSI-H) phenotypes (7–12), while the BLM helicase has gained traction as a drug target to sensitize resistant cancers to alkylating drugs and PARP inhibitors (13–16). Similarly, RECQL4 has entered the spotlight as a tentative druggable target to complement existing cancer therapies (13,17,18).

Despite its established roles in genome integrity and potential status as a prognostic marker for osteosarcoma, breast, and prostate cancers, the physiological functions of RECQL4 remain poorly understood compared to WRN and BLM (13,18–21). This can be partially attributed to the lethality associated with its loss. RECQL4 is an evolutionary chimera, with an N-terminal domain functioning similar to the essential replication initiation factor Sld2 in lower eukaryotes (22) fused to a C-terminal RecQ helicase. Though a majority of its disease-linked and cancer-associated mutations are in the helicase and conserved C-terminal domains (23), RECQL4 helicase activity is not vital to replication (24). However, the *in vivo* study of these alleles remains difficult due to the confounding and potentially lethal pleiotropic effects of perturbing replication initiation. Additionally, RECQL4 is difficult to study biochemically because its large size (135kDa) and long natively disordered N-terminus (first 450 residues) cause it to overexpress poorly with low solubility and a high risk of aggregation in recombinant protein production hosts (25,26). To circumvent these technical limitations and decouple the protein’s dual functions, the *Saccharomyces cerevisiae* homolog Hrq1 (27) has emerged as a powerful model system for characterizing the biochemical and genetic functions of human RECQL4.

Extensive foundational work has validated *S. cerevisiae* Hrq1 as a structural and functional homolog of RECQL4 (25,28–37). Both proteins are 3’ to 5’ helicases containing the conserved RecQ helicase domain but also have degenerate RecQ C-terminal (RQC) and RecQ4 subfamily-specific RHCD (RecQ4/Hrq1-conserved) domains (27). Fibroblasts from patients with Rothmund-Thomson syndrome, an autosomal recessive disease caused by biallelic mutation of *RECQL4*, are sensitized to chemicals that cause DNA inter-strand crosslinks (ICLs) (38), and overexpression of RECQL4 in gastric cancer cells confers resistance to the ICL-inducing agent cisplatin (39). Similarly, *S. cerevisiae* strains lacking Hrq1 (*hrq1*Δ) are sensitive to ICL damaging agents (25,29,30,36,37,40), suggesting the importance of both proteins in ICL repair. Hrq1 functions in a repair pathway with the nuclease Pso2, a homolog of human SNM1A, and both RECQL4 and Hrq1 can specifically stimulate Pso2 nuclease activity *in vitro* (25,29).

Although Hrq1 helicase activity is required for efficient ICL repair, previous work has revealed an intriguing paradox: cells expressing a catalytically inactive *hrq1-K318A* allele are more sensitive to ICL damage than cells lacking *HRQ1* entirely (25,29,30,36). This observation suggests that Hrq1-K318A does not simply represent a loss-of-function mutation but, instead, interferes with ICL repair in a dominant-negative manner. The mechanistic basis for this interference, however, remains unclear. In particular, it remains unclear which properties of Hrq1 – such as DNA binding, protein stability, or interactions with repair factors – contribute to this dominant-negative phenotype.

Dominant-negative alleles have long served as powerful tools for dissecting the functional architecture of multiprotein complexes and multistep enzymatic pathways (41–44). In DNA repair systems, dominant-negative proteins can reveal rate-limiting steps, obligate interactions, and vulnerabilities that are not apparent from null mutations alone (45–48). For helicases, dominant-negative effects are often attributed to stable DNA binding in the absence of productive remodelling, although direct genetic and biochemical evidence remains limited (49–51).

Here, we used a combination of genetic suppressor screening, structural modelling, and biochemical analyses to define the basis of *hrq1-K318A*-mediated ICL repair defects. By isolating spontaneous suppressors of *hrq1-K318A* sensitivity to the ICL-inducing compound diepoxybutane (DEB), we identified intragenic mutations that alleviate dominant-negative toxicity. Computational modelling and functional characterization of a subset of these suppressors revealed that disruption of Hrq1 DNA binding – either through destabilization of the protein or through targeted impairment of DNA interaction – is sufficient to suppress the ICL repair defect caused by catalytic inactivation. Together, these findings support a model in which DNA engagement by a catalytically inactive RecQ4-family helicase contributes to dominant-negative interference with ICL repair.

## MATERIALS AND METHODS

### Chemical reagents

DEB was purchased from Sigma-Aldrich (St. Louis, MO, USA), dissolved in anhydrous DMSO to 1 g/mL, and stored at 4°C in the dark. Working concentrations of DEB were produced by further dilution in DMSO or direct dilution into yeast media. ATP was from DOT Scientific (Burton, MI, USA). All other reagents were molecular biology grade or the highest purity available.

### Yeast strains

All *S. cerevisiae* strains used in this work (Supplementary Table S1) were derivatives of YPH499 (*MATa ura3-52 lys2-801_amber ade2-101_ochre trp1*Δ*63 his3*Δ*200 leu2*Δ*1*) (52). Genes were deleted as described in (53); the details of strain construction are available upon request. All yeast cells were grown in standard media at 30°C with aeration unless otherwise noted. The yeast expression plasmids used in this work are listed in Supplementary Table S2.

### Spot dilution assay

Spot assays were used to compare the sensitivity of wild-type and mutant *S. cerevisiae* strains to the ICL-inducing agent DEB. Generally, 5-mL YPD cultures of indicated strains were incubated overnight at 30°C on a roller drum. Then, the optical density at 600 nm (OD_600_) of each stain was measured using a BioPhotometer spectrophotometer (Eppendorf; Hamburg, Germany), and the strains were diluted to an OD_600_ of 1.0 in 100 μL sterile water in a 96-well plate. Each strain was serially diluted tenfold four additional times, and 5 μL of each dilution was then spotted onto a room temperature YPD or YPD + DEB plate in a grid pattern. Plates were grown at 30°C for 2-3 days and imaged with a Bio-Rad (Hercules, CA, USA) ChemiDoc imaging system or a flat-bed scanner. The YPD + DEB plates were made by supplementing freshly autoclaved and slightly cooled (∼65°C) YPD medium with DEB to the indicated final concentration, mixing thoroughly, pouring into Petri plates, and curing overnight. Because DEB is highly reactive, the plates were typically used within 24 h.

### Spontaneous suppressor screen

To collect spontaneous suppressors of sensitivity to DEB, 100 μL of OD_600_ = 0.01 *S. cerevisiae* culture was plated onto YPD media containing 250 μg/mL (wild-type), 130 μg/mL (*hrq1Δ*), 70 μg/mL (*hrq1-K318A*), or 50 μg/mL (*pso2Δ*) DEB. After 2 d of growth at 30°C, potential suppressor colonies were identified as colonies growing noticeably better than the background growth visible on the plate. These putative suppressors were restreaked onto YPD plates and YPD containing the same concentration of DEB used initially, with their growth compared to a streak of the parental strain. Any clone displaying better growth than the parental strain on YPD + DEB was then evaluated in a spot dilution assay to semi-quantitatively gauge the level of suppression of DEB sensitivity. Because DEB is a mutagen, reiterative exposure to it during the screening process above would result in the accumulation of mutations at each step. Thus, colonies (that were ultimately proven to be suppressors at the spot dilution stage) from the initial YPD restreak were used for subsequent steps.

Genomic DNA (gDNA) from suppressors was harvested from small YPD overnight cultures using a MasterPure Yeast DNA Purification kit (Lucigen; Middleton, WI, USA), following the manufacturer’s instructions. The gDNA was then submitted to the Indiana University Center for Genomics and Bioinformatics (CGB) for small genome sequencing. Libraries were generated with a Nextera Small Genome DNA Library kit and sequenced using a NextSeq instrument (Illumina; San Diego, CA, USA). Bioinformatics support (read mapping and visualization) was also supplied by the CGB.

### Amplicon sequencing of the *hrq1-K318A* locus

To sequence just the *hrq1* locus in *hrq1-K318A* suppressor clones, dozens of suppressors were generated and their gDNA was prepared as described above. Then, the locus was amplified by PCR using primers MB527 (5⍰-GTGAATTGCTCAGAAGAGAAAGGCATACCGTC-3⍰) and MB528 (5⍰-CTGTGCATCAACAAGGTGACAGAATGTTGATG-3⍰) and Bio-Rad iProof HF Master Mix. The DNA was purified using a GeneJet PCR Purification kit (Thermo Fisher Scientific; Waltham, MA, USA), and amplicon sequencing was performed by Plasmidsaurus (Louisville, KY, USA). The suppressor *hrq1* sequences were aligned with the *hrq1-K318A* sequence using CLUSTAL W (54) to identify mutations.

### Western blotting

Yeast protein extracts were prepared by adapting the von der Haar hot alkaline lysis procedure (55). Briefly, *S. cerevisiae* strains containing 3xFLAG-tagged wild-type or mutant Hrq1 (Supplementary Table S1) were grown in 5 mL of YPD overnight. The OD_600_ of each strain was measured, and the cultures were diluted to OD_600_ = 0.09 in 50 mL of 30°C YPD grown to an OD_600_ = 1.0. The equivalent of 9 ODs of culture was harvested by centrifugation (4000xg, 5 min) and then resuspended in freshly prepared protein extraction buffer (0.1 M NaOH, 0.05 M EDTA, 2% SDS, and 2% β-mercaptoethanol). The samples were then heated at 95°C for 10 min before being neutralized with 5 µL of 4 M acetic acid. Fifty microliters of loading buffer (250 mM Tris-HCl [pH 6.8], 8% SDS, 40% glycerol, 0.02% Bromophenol Blue, and 200 mM DTT) was added to each sample, and the lysates were then clarified by centrifugation at 14,000 rpm for 5 min in a microcentrifuge. A small aliquot (10 μL) of each sample was loaded onto a 4-15% SDS-PAGE gel and separated at 140 V for 50 min.

After electrophoresis, the gels were removed from their glass plates and gently rinsed with deionized water. Proteins were transferred to nitrocellulose membranes in transfer buffer (25 mM Tris-HCl [pH 8], 192 mM glycine, and 20 mM methanol) at 400 mA for 80 min at 4°C. After transfer, membranes were cut in half at the BLUEstain 2 protein ladder 75 kDa marker (Gold Biotechnology; St. Louis, MO, USA), and each half was blocked in TBS-T (25 mM Tris-HCl [pH 8], 125 mM NaCl, and 0.1% Tween-20) containing 5% (w/v) non-fat dry milk for 30 min at room temperature with gentle rocking. After blocking, the buffer was replaced with fresh TBS-T/5% non-fat dry milk containing a 1:2000 dilution of primary antibody (top half: monoclonal ANTI-FLAG M2-peroxidase antibody [Sigma, A8592]; bottom half: GAPDH loading control antibody [Invitrogen, MA5-15738]) and rocked overnight at 4°C. The next day, membranes were washed five times with 20 mL TBS-T for 5 min each wash. TBS-T containing a 1:15,000 dilution of the secondary antibody (IRDYE 800CW goat anti-mouse IgG secondary antibody, LICORbio; Lincoln, NE, USA) was added to the membranes and rocked at room temperature for 30 min. Then, the membranes were washed as above and briefly washed a final time with deionized water. Membranes were scanned on a LICORbio Odyssey DLx, and Hrq1 and GAPDH protein bands were quantified with Image Studio software.

### Pathogenicity and stability predictions

To assess the evolutionary and structural impact of Hrq1 variants, we utilized the ESM-1v protein language model (56). Unlike traditional methods that rely on explicit multiple sequence alignments, ESM-1v is an unsupervised 650-million-parameter transformer model trained on the UniRef90 database. It predicts the functional effect of mutations by calculating the log-likelihood ratio between the mutant and WT amino acids. For the 1077-aa Hrq1 sequence, we implemented a sliding-window approach (1022-aa windows) to accommodate the model’s 1024-token context limit. Mutations were classified as “Highly Deleterious” (score < -10), “Likely Deleterious” (score between -6 and -10), or “Likely Benign” (score > -6) based on established benchmarks.

### Computational Structural Modelling and Energetics

Structural analysis of *S. cerevisiae* Hrq1 was performed using the AlphaFold2-predicted structure (UniProt: Q05549) (57). To prepare the model for thermodynamic calculations, the structure was first processed using the RepairPDB command in FoldX 4.0 (58) to resolve steric clashes and optimize side-chain rotamers. The energetic impact of each identified suppressor mutation was calculated using the BuildModel command. Each mutation was modelled five times to ensure convergence, and the change in Gibbs free energy (ΔΔG) was reported as the average difference between the mutant and wild-type (WT) proteins. Mutations resulting in a ΔΔG > 1.6 kcal/mol were classified as significantly destabilizing, while those exceeding 5.0 kcal/mol were defined as “Structural Kills” likely to result in proteostatic clearance. Complementary stability predictions were performed using MAESTRO (Multi-Agent Stability TRansfOrmation) to provide an empirical consensus of thermodynamic disruption based on statistical potentials (59).

### Protein conservation analysis

To characterize the evolutionary relationship between the *S. cerevisiae* Hrq1, human RECQL4, and *Bacillus subtilis* MrfA RecQ4-family helicases, we retrieved their amino acid sequences from GenBank (accession numbers WNF20109.1, BAA86899.1, and WP_015383941.1, respectfully). The sequences were aligned using CLUSTAL Omega (v1.2.4) (60) to visualize conserved motifs. To characterize the pairwise relationship between these orthologs, both full-length global and catalytic core local alignments were performed using the EMBOSS suite (EBLOSUM62 matrix; Gap Open: 10.0; Gap Extend: 0.5) (61). Global alignments (Needleman-Wunsch via EMBOSS Needle) reflect the total sequence divergence, including the extensive, non-conserved N- and C-terminal extensions found in the eukaryotic sequences. Local alignments (Smith-Waterman via EMBOSS Water) were utilized to isolate and quantify the conservation of the core helicase machinery.

### *In silico* structural modelling

All structural predictions of Hrq1 (Supplementary File S1) were rendered using AlphaFold on the publicly accessible Google DeepMind AlphaFold server (https://alphafoldserver.com/). Structures were rendered with the default parameters, and figures display the overall highest-confidence model. The wild-type Hrq1 protein sequence used for the modelling is from the S288c genotype (https://yeastgenome.org/locus/S000002699), and all Hrq1 mutants were constructed using that template. The *Bacillus subtilis* MrfA structure (62) was retrieved from the protein Data Bank (https://www.rcsb.org/structure/6ZNS). Sequences were visualized in either Chimera X (https://www.cgl.ucsf.edu/chimerax/) or PyMOL (https://www.pymol.org/).

### Baculovirus production and insect cell infection

Baculovirus transfer vectors encoding *HRQ1* or mutant *hrq1* genes were constructed as described in (30). Details of plasmid construction are available upon request. Bacmid transfection was performed using Cellfectin II (Gibco; Waltham, MA, USA) and following the manufacturer’s protocol. Briefly, 3 µg of purified bacmid DNA was diluted into 100 µL of Hyclone SFX-insect cell culture medium not containing antibiotics. This dilution was combined with 8 µL of Cellfectin II diluted into 100 µL of the same media, gently flicked to mix, and incubated at room temperature for 15-30 min. During this incubation, 8×10^5^ cells of a low-passage Sf9 insect cell culture was plated onto a 6-well culture plate and left stationary at room temperature for 15 min to allow cells to adhere to the wells. Subsequently, all media was gently aspirated from the wells and replaced with 2 mL of antibiotic-free SFX medium. The roughly 210-µL bacmid-Cellfectin II mixture was then added dropwise to the well. The edges of the 6-well plate were wrapped in a layer of parafilm and incubated at 27°C for 3-5 h. The transfection medium was then aspirated and replaced with 2 mL of SFX medium supplemented with a mix of penicillin, streptomycin, and Amphotericin B (Gibco; Waltham, MA, USA). The plate was again wrapped with parafilm and incubated at 27°C for 4 d. The viral supernatants were harvested from the wells and immediately used for a round of infection.

To amplify the virus further, 20 µL of the viral supernatant harvested from transfection was used to infect 2×10^6^ Sf9 cells in 2 mL of SFX media containing antibiotics in a fresh 6-well plate. This plate was wrapped with parafilm and incubated at 27°C for 4 d. Viral supernatants were harvested from the wells and stored in sterile microcentrifuge tubes at 4°C for up to 2 weeks. Optionally, an additional, identical round of infection can be done with the virus from the first round of infection to further increase the yield of high-titre virus, though this was typically not required. For protein expression, 500 mL Sf9 culture at a density of 1.5×10^6^ cells/mL and >96% live count was infected with 500 µL of high-titre virus (1:1000 dilution)and incubated at 27°C with shaking at 140rpm. Infected cells were harvested after the live count percentage dropped to 70-80% (approximately 48-60 h) by centrifugation at 2000xg for 20 min and stored at -80°C prior to protein purification.

### Purification of Hrq1 and Hrq1 mutants

The following protocol was adapted from (28,30). Briefly, frozen pellets from 500 mL infected Sf9 cultures were thawed on ice and resuspended in 200 mL of ice-cold Hrq1 lysis buffer (50 mM Na-HEPES [pH 7.6], 0.4 M NaCl, 20 mM imidazole, and 10% glycerol) supplemented on the same day with protease inhibitor mix (600 nM leupeptin, 2 μM pepstatin A, 2 mM benzamidine, and 1 mM phenylmethanesulfonyl fluoride [PMSF]), 20 μg/ml DNase I, and fresh 5 mM β-mercaptoethanol. The resuspended cell pellet was further lysed via 10 strokes with a Dounce homogenizer on ice. Lysate was clarified by centrifugation at 4°C for 30 min at 14,000 rpm and then loaded onto a lysis buffer-equilibrated gravity column containing 3 mL of HIS-Select Nickel Affinity Gel (Sigma) at 4°C. The resin was washed once with 15 mL lysis buffer containing 5 mM ATP and then twice more with 30 mL of unadulterated lysis buffer. Hrq1-6xHis was eluted using three column volumes of Hrq1 lysis buffer supplemented with 500 mM imidazole. Fractions containing Hrq1 (determined by SDS-PAGE and Coomassie blue staining) were pooled and concentrated to < 1 mL using a 30 kDa MWCO Amicon centrifugal filter at 4°C. The concentrated protein sample was then injected onto a Superdex 200 10/300 GL gel filtration column with a Bio-Rad NGC FPLC system that was pre-equilibrated with Hrq1 storage buffer (25 mM Na-HEPES [pH 8], 30% glycerol, 300 mM NaOAc [pH 7.6], 25 mM NaCl, 5 mM MgOAc, 1 mM DTT, and 0.1% Tween-20). Hrq1 was eluted with storage buffer at a rate of 0.1 mL/min and collected in 500-µL fractions. Fractions containing Hrq1 were identified using SDS-PAGE and Coomassie blue staining, pooled, and concentrated as above. Concentrated, purified Hrq1 was dispensed into 15-µL aliquots and flash frozen in liquid nitrogen before being stored at -80°C. All Hrq1 mutants were purified identically.

### Quantification of recombinant proteins

Our staining procedure was adapted from (63,64). Briefly, a colloidal solution of Coomassie blue G-250 was made by dissolving 100 g of anhydrous ammonium sulphate in 650 mL of deionized water in a 1-L graduated cylinder, stirring until fully dissolved. Then, 100 mL of 200-proof ethanol was added and mixed thoroughly, followed by 200 mg of Coomassie Brilliant Blue G-250 powder, which was stirred until completely dispersed (∼20 min). Finally, 10 mL of 85% phosphoric acid was added dropwise while stirring, and then the graduated cylinder was filled to a final volume of 1 L with deionized water. The stain was mixed on a stir plate for ≥ 30 min before use.

Purified proteins were loaded onto 4-15% gradient SDS-PAGE gels alongside BSA standards and were run at 140 V for 50 min. The gels were then removed from their glass plates and soaked in deionized water on a rocker for 30 min. The rinsed gels were placed into clean containers with colloidal Coomassie G-250 stain and rocked for > 2 h. The gels were removed from the stain and quickly rinsed in a clean container several times with deionized water. Then, the gels were rocked in deionized water for > 2 h to destain. Gels were imaged on a Bio-Rad ChemiDoc imaging system, and bands were quantified using ImageJ.

### Mass photometry

Mass photometry was used to determine the molecular masses of recombinant proteins and their in-solution oligomerization states, essentially as described (65,66). All analyses were performed on a Refeyn TwoMP system (Oxford, UK) using the accompanying software AcquireMP and DiscoverMP for data acquisition and analysis, respectively. All proteins were diluted to a final concentration of 20 nM in freshly prepared PBS buffer (filtered through a 0.45-µM PES membrane filter and then through a 0.1-µM MCE membrane filter [Millipore; Burlington, MA, USA]). Recordings for each protein were collected over 180 s, and the readings were calibrated with the MassFerence P1 calibration standard (Refeyn) diluted into the same PBS buffer and recorded within 1 h of the other protein measurements.

### Electrophoretic mobility-shift assay (EMSA)

EMSAS were performed to measure the DNA binding activity of the purified helicases. Protein was incubated at the indicated concentrations with 1 nM of a single-stranded poly(dT)_30_ oligonucleotide containing a 5⍰ IRDye700 label (MB1755; 5⍰-/5IRD700/TTTTTTTTTTTTTTTTTTTTTTTTTTTTTT-3⍰) for 30 min at 30°C in DNA binding buffer (25 mM Na-HEPES [pH 8.0], 5% glycerol, 50 mM NaOAc [pH 7.6], 150 μM NaCl, 7.5 mM MgOAc, and 0.01% Tween-20). Free single-stranded DNA (ssDNA) was separated from ssDNA-protein complexes on 8% 29:1 acrylamide:bis-acrylamide native-PAGE gels in TG buffer at 100 V for 50 min at 4°C. Gels were imaged using a LICORbio DLx imager, and bands were quantified with ImageStudio software.

### Helicase assays

The DNA unwinding activity of wild-type and mutant Hrq1 was measured using 200 nM of the indicated protein and 1 nM forked DNA substrate. The probe was constructed by incubating 10 μM oligonucleotide MB1772 (5⍰-/5IRD700/GAACGCTATGTGAGTGACACTTTTTTTTTTTTTTTTTTTTTTTTT-3⍰) with 11 μM MB733 (5⍰-ACCGTTGTGCAACTGAGTGGACAACGTGTCACTCACATAGCGTTC-3⍰) in a 95°C heat block for 5 min in annealing buffer (20 mM Tris-HCl [pH 8], 4% glycerol, 0.1 mM EDTA, 40 µg/mL BSA, 10 mM DTT, and 10 mM MgOAc). The heat block was then unplugged, and the substrate was allowed to slowly cool to room temperature over 3 h. This created a fork substrate with a 20-bp duplex region, a 25-nt random-sequence 5⍰-tail, and a 25-nt poly(dT) 3⍰-tail with nearly 100% efficiency. For the helicase assays, protein was incubated with labelled fork and 100 nM unlabelled MB1772 for 30 min at 30°C in DNA binding buffer containing 2 mM ATP. The reactions were stopped with an equal volume of 2x Proteinase K/SDS stop buffer (20 mM EDTA [pH 8], 1% SDS, 20% glycerol, 0.1% Orange G Dye, and 1 mg/mL Proteinase K). Unwound substrate was separated from annealed substrate, imaged, and quantified as described for the EMSAs above.

### Statistical analyses

Data were analysed and graphed using GraphPad Prism software. The reported values are averages of ≥ 3 independent experiments, and the error bars are the standard deviation. *P*-values were calculated as described in the figure legends, and we defined statistical significance as *p* < 0.01.

## RESULTS

### A spontaneous suppressor screen for DEB ICL repair factors

We previously reported that the transcription of three genes of unknown function - *FMP48*, *TDA6*, and *YLR297W* - is significantly upregulated in cells expressing the *hrq1-K318A* allele (34). We hypothesized that perhaps these genes encode proteins involved in DNA ICL repair and that yeast cells upregulated them in the context of a crippled Hrq1-Pso2 ICL repair pathway to compensate for its lack of activity (34,67). We tested these candidate genes for roles in DEB ICL repair but did not observe sensitivity in the deletion strains alone or in combination with disruption of the Hrq1-Pso2 (*hrq1Δ*) or proto-Fanconi anaemia (FA; *chl1Δ*) (30) repair pathways (Fig. S1). Thus, these epistasis analyses suggest that the *FMP48*, *TDA6*, and *YLR297W* gene products do not participate in DEB ICL repair in yeast, though they may be involved in the repair of ICL lesions formed by other compounds such as mitomycin C or cisplatin.

As an alternate method to identify *S. cerevisiae* proteins involved in DEB ICL repair, we performed a spontaneous suppressor screen for mutants displaying resistance to DEB. The basis of this screen was the observation that strains exposed to lethal doses of DEB occasionally yield colonies that are resistant to the concentration used (Fig. 1A). We took advantage of this phenomenon by performing a screen as depicted in Figure 1B. Briefly, *S. cerevisiae* strains were exposed to varying doses of DEB (higher for resistant strains like WT, lower for sensitive mutants like *pso2Δ*) on plates, and the suppression in ICL sensitivity of colonies that arose was tested using spot dilution assays on media containing DEB and compared to the parental strain. Genomic DNA was then harvested from verified suppressors and subjected to whole-genome sequencing to identify the genetic source of the resistance to ICL damage. Multiple rounds of the initial plating were used to avoid jackpot mutations (68) that could bias the results. We hypothesized that we would identify mutations that altered the cell wall or cell membranes such that DEB entry into the cells was decreased, mutations in drug pumps that increased their ability to evict DEB from the cells, and mutations that activate or upregulate DNA repair proteins. Importantly, this approach enabled the identification of mutations that suppress the *hrq1-K318A* dominant-negative phenotype, irrespective of whether suppression occurs through restoration of repair activity or through alleviation of toxic interference.

**Figure 1.**
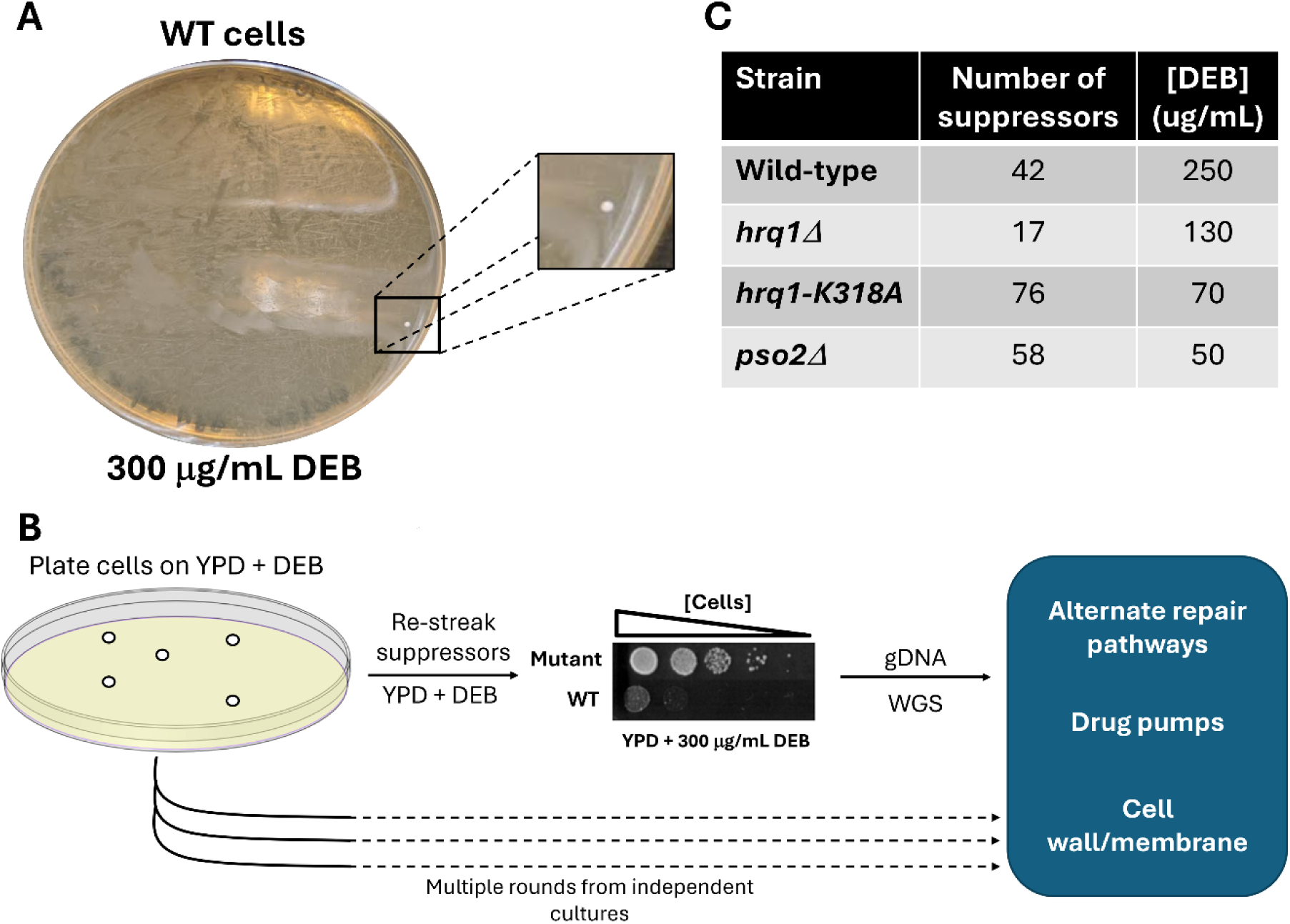
Screen for suppressors of DEB sensitivity. **A)** Spontaneous suppressor of DEB sensitivity. WT cells were plated on rich medium containing a lethal dose of DEB, but a single suppressor mutant arose and generated a colony. **B)** Schematic of the suppressor screen. Briefly, suppressors will be collected as in (**A**), restreaked to verify resistance to DEB, compared to the parental strain using spot dilution assays, and then genomic DNA (gDNA) will be harvested for whole-genome sequencing (WGS). Putative pathways affected by suppressor mutations are shown. **C)** Average number of suppressors *vs.* DEB concentration. The indicated strains were grown, in triplicate, on rich media supplemented with the indicated concentrations of DEB, and the average number of suppressor colonies recovered is reported.

Initially, we performed this screen in *hrq1* and *pso2* mutant backgrounds in an attempt to identify factors that can compensate for the loss of the main DEB ICL repair pathway. However, we were also able to gather spontaneous suppressors of DEB sensitivity in WT cells in hopes of gaining a more global view of repair. By adjusting the concentration of DEB used to fit the sensitivities of the individual strains analysed, we were able to recover dozens of suppressors per strain (Fig. 1C). Overall, we sequenced the genomes of approximately 8-10 suppressors from WT, *hrq1Δ*, and *hrq1-K318A* strains, as well as one pilot suppressor from the *pso2Δ* background (Table 1). The average number of mutations identified per genome roughly scaled with the basal sensitivity of the strains to ICL damage, with the fewest found in WT suppressors and the most in the single *pso2Δ* suppressor. Our initial analyses of these genomics data revealed few common threads, and a technique like bulk segregant analysis (69) will be necessary to identify the genetic cause of suppression for each clone. As that work is ongoing, we instead focus here on suppressors in the *hrq1-K318A* background, which all shared one commonality described below.

**Table 1.** Whole-genome sequencing results.

| Table 1. Whole-genome sequencing results |  |  |  |
| --- | --- | --- | --- |
| Strain | No. suppressors screened | Average mutations per genome | Relative sensitivity to DEB* |
| WT | 9 | 27 | + |
| *hrq1Δ* | 10 | 67 | ++ |
| *hrq1-K318A* | 8 | 85 | +++ |
| *pso2Δ* | 1 | 100 | ++++ |

\*Relative sensitivity: WT is the least sensitive, and *pso2Δ* is the most sensitive to DEB-induced ICL damage.

### Suppressors of *hrq1-K318A* ICL sensitivity are secondary mutations in the helicase

The eight suppressor clones in the *hrq1-K318A* strain contained an average of 85 DEB-induced mutations (Table 1). Strikingly, all suppressors identified in the *hrq1-K318A* background contained secondary mutations within *HRQ1* itself (Fig. 2A). This finding indicates that in this genetic background, suppression primarily occurs through modification of the mutant protein rather than through activation of compensatory pathways.

**Figure 2.**
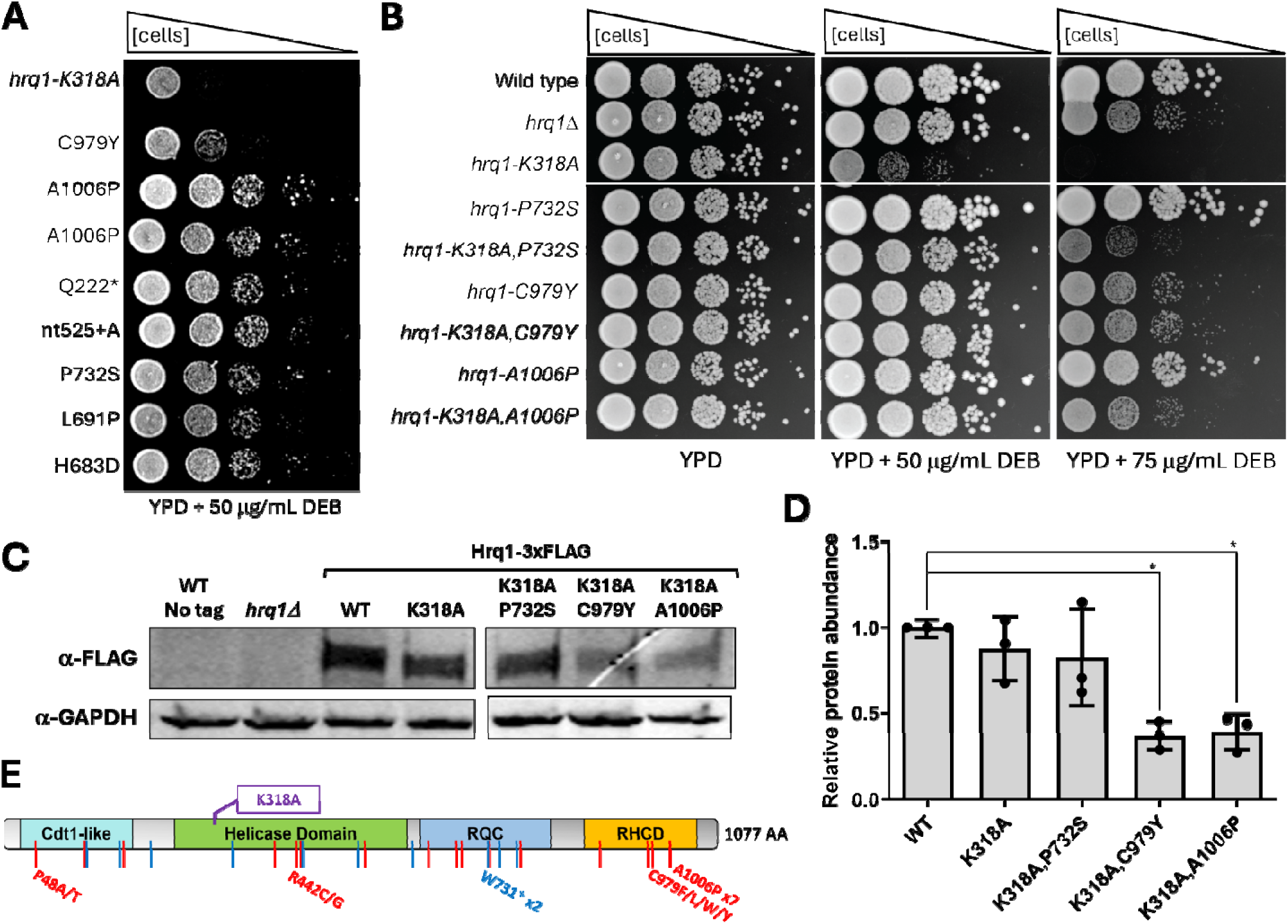
Second-site mutations in *hrq1-K318A* suppress ICL sensitivity. **A)** Spontaneous suppressor clones from our screen contain secondary mutations in the *hrq1-K318A* locus. Ten-fold serial dilutions of *hrq1-K318A* suppressors containing the mutation listed on the left are shown. The Q222* mutation denotes that Q222 was altered to a stop codon in this strain. The nt525+A mutation denotes the addition of an adenine at nucleotide position 525 in the *hrq1* gene, resulting in a frameshift and early termination in the protein sequence. **B)** Second-site mutations were remade in both wild-type cells the parental *hrq1-K318A* strain, and their sensitivity to DEB was compared to wild-type, *hrq1Δ*, and the *hrq1-K318A* allele alone. **C)** The selected suppressors are produced as full-length proteins *in vivo*. Western blot analysis of protein extracts from the indicated strains. GAPDH protein levels were used as a loading control. **D)** Quantification of n = 3 blots as performed in (C). 3xFLAG-tagged protein abundance was normalized to GAPDH levels in each experiment and then further normalized to wild-type (WT) Hrq1. The amount of Hrq1-K318A,C979Y and Hrq1-K318A,A1006P protein was significantly reduced compared to WT as determined by one-way ANOVA. * *p* < 0.01. **E)** Domain schematic of Hrq1 showing the sites of the suppressor mutants recovered, as well as the location of the K318A mutation. Red hashes denote substitution mutations, and blue lines denote frameshifts/premature stop codons. RQC, RecQ C-terminal domain; RHCD, RecQ4/Hrq1-conserved domain.

With the exception of the A1006P mutation, which was independently collected twice, all of these second-site mutations were unique, with some causing amino acid substitutions and others introducing frameshifts and/or premature stop codons. To determine if these mutations were responsible for the resistance to DEB, we remade several of them in the parental *hrq1-K318A* strain, yielding cells with two mutations in *hrq1* in an otherwise WT background. In all cases, when plated on medium containing DEB, the double-mutant strains displayed resistance compared to the single *hrq1-K318A* mutant (Fig. 2B and S2). This indicates that the suppression of sensitivity to DEB is a monogenic trait and not due to the cumulative action of multiple mutations in these suppressor clones. Western blot analysis of several of the full-length mutants confirmed that they are produced *in vivo* (Fig. 2C and S3), though the protein levels of the Hrq1-K318A,C979Y and Hrq1-K318A,A1006P mutants were significantly reduced (*p* = 0.0036 and 0.0049, respectively) compared to WT Hrq1 (Fig. 2D).

It should be noted that cells expressing the *hrq1-K318A* allele are more sensitive to ICL damage than *hrq1Δ* cells (Fig. 2B). This phenotype has been observed before (29,30,70), and we hypothesized that it is due to the *hrq1-K318A* allele being a dominant negative. The protein is expressed at WT levels (Fig. 2D), and the model is that it is recruited to its sites of action at ICL damage (29). However, because it is catalytically inactive, the Hrq1-K318A protein then serves as a roadblock to compensatory repair mechanisms that may be able to repair the lesions. To determine if *hrq1-K318A* is a dominant negative, we cloned *HRQ1* under the control of its native promoter in a single-copy vector, transformed it into *hrq1Δ* and *hrq1-K318A* cells, and then tested sensitivity to DEB relative to cells containing empty vector (Fig. S4). Although *HRQ1* expression was able to rescue the DEB sensitivity of *hrq1Δ* cells, it was much less effective in the *hrq1-K318A* background, supporting the notion that the *hrq1-K318A* allele is dominant negative.

### Suppressor mutations occur in conserved residues

Because the suppressor screen performed in the *hrq1-K318A* strain yielded 100% second-site suppressors in the *hrq1-K318A* gene itself, we reasoned that sequencing just the *hrq1* locus in additional suppressors collected in this genetic background would reveal how common this route of suppression is. To do this, we analysed dozens of additional DEB sensitivity suppressors from the *hrq1-K318A* strain collected from the screen depicted in Figure 1B and amplified the *hrq1* locus and flanking regions (∼4 kb) for sequencing. From 43 total suppressors, 36 (84%) contained a second-site mutation in the *hrq1-K318A* gene (Table 2). Although mutations at some residues (*i.e.*, P48, R442, W731, C979, and A1006) were recovered more than once, the majority were unique. In some cases, more than one additional mutation was found in the *hrq1-K318A* gene, but such events were less common. Figure 2E shows the sites of these mutations along a domain schematic of Hrq1, revealing no apparent hotspots but, rather, that suppression of DEB sensitivity can result from mutating any of the domains in Hrq1.

**Table 2.** Hrq1-K318A second-site suppressors and the conservation of the mutated residues in *Bacillus subtilis* MrfA and/or human RECQL4.

| Additional Hrq1-K318A mutation(s) | No. times identified | Conserved residue in MrfA | Conserved residue in RECQL4 | RECQL4 mutation in COSMIC database |
| --- | --- | --- | --- | --- |
| P48A | 1 | - | P199 | P199S |
| P48T | 1 | - | P199 | P199S |
| Q121K , Q125* | 1 | N/A | N/A | P285Hfs*8 |
| N122Y | 1 | - | S251 | S251I, S251R |
| nt525+A | 1 | N/A | N/A | N/A |
| F181L , V641G | 1 | - , V397 | F342 , F817 | - , - |
| Q222* | 1 | N/A | N/A | G392Vfs*14 |
| nt1035,1036-GG | 1 | N/A | N/A | C525Afs*33 |
| N409K | 1 | K171 | Q595 | Q595P |
| R442C | 1 | R204 | - | N/A |
| R442G | 1 | R204 | - | N/A |
| H448N , Y451* | 1 | N/A | N/A | N/A |
| nt1607+T | 1 | N/A | N/A | - |
| L545F | 1 | I304 | I716 | - |
| nt1852+A | 1 | N/A | N/A | A805Pfs*38 |
| H683D | 1 | H435 | - | N/A |
| L691P | 1 | L443 | - | N/A |
| W731* | 2 | N/A | N/A | W934* |
| P732S | 1 | P484 | - | N/A |
| nt2239-G (A747P , V748*) | 1 | N/A | N/A | Q957* |
| nt2322-C | 1 | N/A | N/A | W998* |
| Q780K , S972N | 1 | E533 , A711 | - , - | N/A |
| A900E | 1 | - | G1118 | G1118S |
| C979F | 1 | C718 | - | N/A |
| C979L | 1 | C718 | - | N/A |
| C979W | 1 | C718 | - | N/A |
| C979Y | 1 | C718 | - | N/A |
| G1005C | 1 | - | - | N/A |
| A1006P | 7 | - | - | N/A |
| None | 7 | N/A | N/A | N/A |

**N.B.** The orange shading indicates different substitutions at the same residue. - indicates that the residue is not conserved and/or not mutated in the COSMIC database. Frameshift and early termination mutations were not analysed for conservation and are listed as not applicable (N/A). Frameshift and early stop codon mutations found in the Catalogue Of Somatic Mutations In Cancer (COSMIC) database were within ~10 aa of the analogous residue in RECQL4.

To determine if the second-site suppressors occurred in conserved residues, we aligned the *S. cerevisiae* Hrq1 sequence (1077 aa) with the human RECQL4 (1208 aa) (30) and *Bacillus subtilis* MrfA (749 aa) (71) helicases (Fig. S5). We chose these specific homologs to define the ‘minimal essential’ RecQ4-family helicase. They represent evolutionarily distant members of the RecQ4 family and allow identification of residues conserved across broad phylogenetic distances. Globally, RECQL4 is 14.6% identical and 27.0% similar to Hrq1, while MrfA is 23.2% identical and 38.3% similar to Hrq1. As these proteins differ considerably in size, largely due to variability in their N- and C-terminal domains (Fig. S5), local alignment of the RecQ core of the enzymes revealed stronger conservation: 24.3% identity and 40.2% similarity between RECQL4 and Hrq1; and 32.1% identity and 53.1% similarity between MrfA and Hrq1.

As shown in Table 2, nearly all of the secondary missense mutations in Hrq1-K318A hit residues that were conserved in RECQL4 and/or MrfA, suggesting that they are important for the inherent function or stability of these RecQ4-family enzymes. Indeed, >60% of the Hrq1 missense mutations that were in conserved in RECQL4 are also sites of RECQL4 variants in the Catalogue Of Somatic Mutations In Cancer (COSMIC) database (72) (Table 2 and Supplementary File S2), which suggests they could be related to the cancer predisposition common of RECQL4-related diseases (3,73,74). The only two suppressor mutations that were not in conserved residues in either RECQL4 or MrfA, G1005C and A1006P, are found in the divergent C-terminal tail of Hrq1 (Fig. S5), which is only found in fungal and plant homologs (27).

We also recovered nonsense mutations in our screen that resulted in frameshifts and early stop codons (Table 2). Similarly, two missense mutations that introduced stop codons (Q222* and W731*) were found. Mutations in these latter two categories were not analysed for conservation due to the large truncations that they result in, which presumably yield inactive or unstable proteins. However, the COSMIC database does include such mutations (72), many of which could be found within ∼10 aa of the analogous residue in RECQL4 (Table 2 and Supplementary File S2).

### Multi-algorithm *in silico* profiling suggests functionally critical mutations

No mutational hotspots are evident along the primary sequence of Hrq1 (Fig. 2E). To predict the evolutionary and structural impact of the Hrq1 mutations identified as suppressors of DEB sensitivity, we performed comprehensive *in silico* analyses using three independent scoring frameworks: ESM-1v (evolutionary fitness; (56)), FoldX (empirical force-field stability; (58)), and MAESTRO (statistical potential stability; (59)). The outputs of these methods are listed in Supplementary Table S3 and visualized as heatmaps in Figure 3A. Notably, these analyses were restricted to the 24 identified missense mutations. Suppressor mutations resulting in frameshifts or premature termination codons (Fig. 2E) were excluded from these modelling pipelines, as the resulting truncated or significantly altered proteins lack the defined tertiary fold required for scoring.

**Figure 3.**
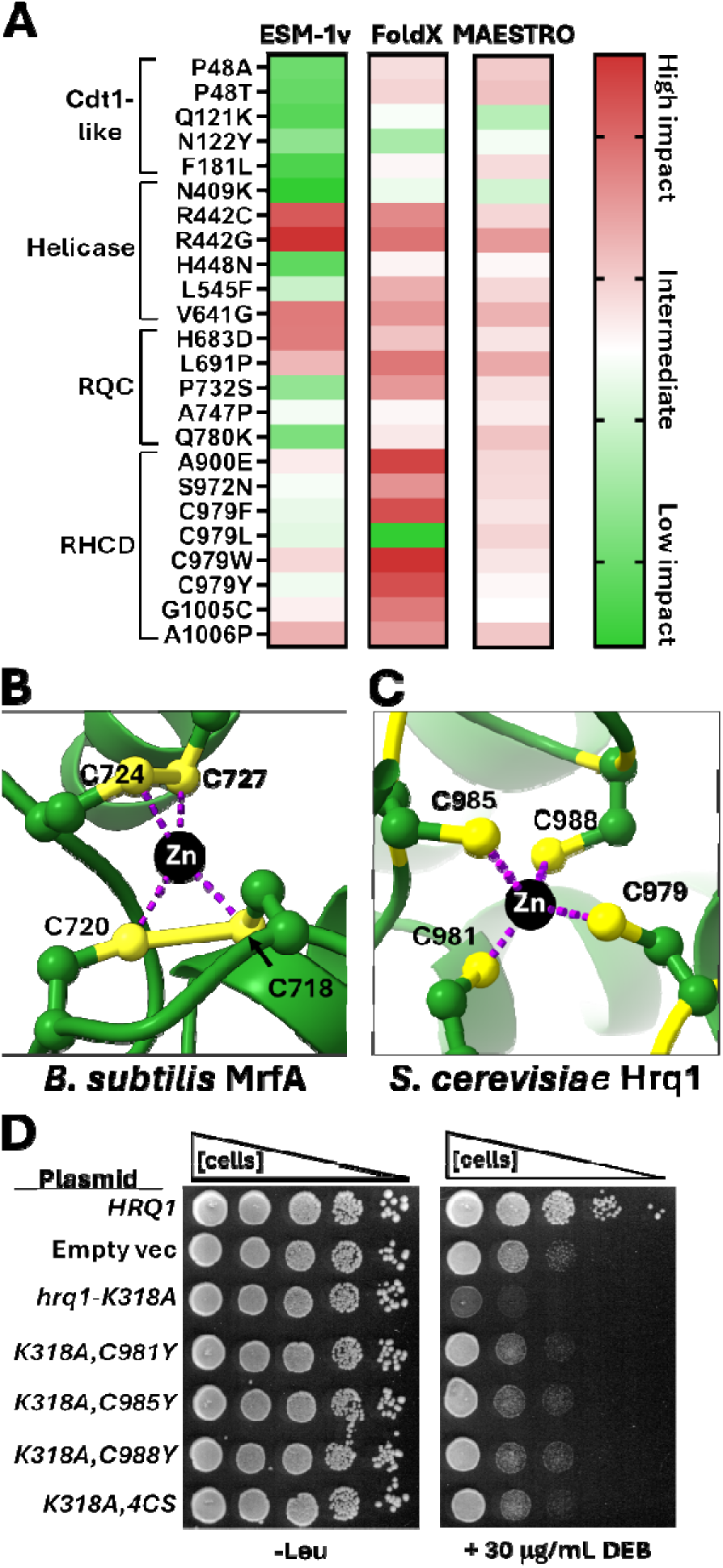
In silico analysis of suppressor mutations. **A)** Heatmaps summarizing the predicted impact of 24 missense suppressor mutations on Hrq1-K318A function and stability. Predictions were generated using three independent computational pipelines: ESM-1v for evolutionary conservation and structural context, FoldX for atomic-scale force field ΔΔG calculations, and MAESTRO for statistical potential-based stability assessment. The heatmaps share a combined colour scheme from low i pact (green = likely benign or neutral) to high impact (red = highly deleterious or destabilizing). **B)** The 4-Cys Zn^2+^ coordination site from the B. subtilis MrfA crystal structure (PDB: 6ZNS). **C)** The predicted str cture of the 4-Cys motif in *S. cerevisiae* Hrq1 with Zn^2+^ modelled in. **D)** Mutating any residue in the 4-Cys motif suppresses the DEB sensitivity of the *hrq1-K318A* mutant.

First, using the protein language model ESM-1v, we identified a subset of mutations with “Highly Deleterious” scores (<−10.0), suggesting these residues are under intense purifying selection. As an internal control, the Walker A box K318A mutation, which catalytically inactivates the helicase (25), was also analysed and yielded a score of -14.939 (Table S3), establishing a baseline for total functional loss. Other highly deleterious mutations included R442G (-12.554), V641G (-10.671), and H683D (-10.608). Conversely, variants such as N409K (-1.729) and Q121K (-2.752) were classified as “Likely Benign,” suggesting they represent tolerated mutations that do not significantly impact the protein’s evolutionary fitness (Fig. 3A, Table S3).

Next, to determine if these evolutionary penalties were due to protein instability, we calculated changes in folding free energy (ΔΔG). This required a three-dimension structure of the enzyme, but no high-resolution structural data exist for Hrq1. Instead, we used AlphaFold to predict the helicase’s structure (Fig. S5 and Supplementary File S1). After analysing the highest-confidence structure with FoldX and MAESTRO, a discrepancy was observed between the two algorithms. While FoldX predicted several mutations to be “Destabilizing” (ΔΔG >1.5 kcal/mol), MAESTRO predictions for all sites remained “Neutral/Marginal” (ΔΔG ≈ -0.5 to 0.5 kcal/mol) (Figure 3A, Table S3). This divergence between evolutionary conservation (ESM-1v) and global structural stability suggests that some of the mutated residues are likely critical for Hrq1 function, potentially coordinating the DNA substrate during the repair of DEB-induced ICLs, rather than being essential for the global fold of the protein.

### The RHCD – a helicase stability module

The integration of evolutionary fitness (ESM-1v) and structural stability (FoldX and MAESTRO) reveals a nuanced landscape of Hrq1 regulation. The degree of consensus, or the lack thereof, between these models provides a roadmap for identifying which mutations disrupt the physical scaffolding of the protein *vs*. those that specifically handicap its biochemical activities. For instance, while most of the mutations in the Cdt1-like domain are predicted to have a low or intermediate impact on Hrq1, there is a striking lack of consensus between ESM-1v and the structural predictors in the catalytic core (helicase and RQC domains) (Fig. 3A). A simple interpretation of this is that the evolutionary model recognizes that these residues are essential for the “engine” of the enzyme, but the structural models suggest that the protein can still fold correctly even when the catalytic “spark plugs” are mutated. This suggests that these suppressors work by modulating the biochemical activity of the helicase rather than by destabilizing the protein. Toward the C-terminus of the protein, there is a higher degree of consensus across the models, with mutations in the RHCD predicted to be of intermediate-to-high impact (Fig. 3A). Indeed, the most deleterious FoldX predictions are found in the RHCD. As this domain is only found in RecQ4-family helicases, we focused on it to investigate how the RHCD may function.

The RHCD of *B. subtilis* MrfA displays a high level of conservation with *S. cerevisiae* Hrq1 (Fig. S5), and a recently reported crystal structure of MrfA (62) shows that a 4-Cys motif coordinates a Zn^2+^ ion (Fig. 3B). We recovered four different suppressor mutations in the Hrq1 residue C979 (Table 2, Fig. 3A, Table S3), which is part of this conserved CXCX_3_CX_2_C motif. We included Zn^2+^ ions in our AlphaFold prediction of the Hrq1 structure (Fig. S5 and Supplementary File S1), which yielded a similar predicted zinc-coordination architecture (Fig. 3C). All four suppressor mutants would disrupt metal ion coordination in this motif, despite ESM-1v and MAESTRO analyses tolerating most of them (Fig. 3A, Table S3). Indeed, FoldX even predicts that C979L to be the most stabilizing of all of the suppressor mutants, demonstrating the limits of these algorithms.

To further probe the Hrq1 4-Cys motif, we individually mutated the other three Cys residues (C981, C985, and C988) to Tyr in the context of the Hrq1-K318A protein to mimic Hrq1-K318A,C979Y. These mutants were cloned into a *LEU2*-marked vector under the control of the *HRQ1* promoter and expressed in *hrq1Δ* cells. As shown in Figure 3D, expressing wild-type Hrq1 with this system rescued the DEB sensitivity of the *hrq1Δ* strain, but Hrq1-K318A expression further sensitized the cells. However, expressing Hrq1-K318A,C981Y, -K318A,C985Y, or -K318A,C988Y phenocopied *hrq1Δ*, as we demonstrated for Hrq1-K318A,C979Y above (Fig. 2B). Previously, more conservative Cys-to-Ser mutations were used to examine the 4-Cys motif in *Mycobacterium smegmatis* RecQ4-family helicase SftH (75), so we also tested a 4xCys-to-Ser Hrq1-K318A mutant (Hrq1-K318A,4CS). Much like the single Cys-to-Tyr mutants, cells expressing Hrq1-K318A,4CS displayed DEB sensitivity like *hrq1Δ* rather than *hrq1-K318A* (Fig. 3D). Thus, mutating any or all of the residues involved in the predicted zinc coordination site suppresses the DEB sensitivity of the *hrq1-K318A* mutation.

Our western blot analysis of Hrq1-K318A,C979Y in Figures 2C and 2D shows a significant reduction of protein abundance relative to WT or Hrq1-K318A, suggesting that the suppressors in the 4-Cys motif function by destabilizing the mutant protein to phenocopy the DEB sensitivity of Hrq1-null cells. This is perhaps unsurprising given that mutation of the homologous CXCX_3_CX_2_C motif in *M. smegmatis* SftH also destabilizes the protein (75).

The most commonly collected *hrq1-K318A* suppressor is the A1006P mutation (Table 2), which is also located in the RHCD but in a portion not conserved with MrfA (Fig. S5). AlphaFold predicts that A1006 resides in a short α-helix (K1003-L1014) (Fig. S6 and S7). Interestingly, α-helix_1003-1014_ is situated near the 4-Cys motif in the predicted Hrq1 structure (Fig. S7). An Ala-to-Pro mutation at residue 1006 would disrupt the N-terminal one-third of α-helix_1003-1014_, potentially altering the conformation of the protein near the 4-Cys motif, and all three *in silico* algorithms predict that this is detrimental (Fig. 3A and Table S3). *In vivo*, Hrq1-K318A,A1006P protein levels are also significantly reduced compared to WT and Hrq1-K318A (Fig. 2C and 2D). Therefore, the RHCD appears to be a domain in RecQ4-family helicases that is important for protein stability, and mutations in the RHCD likely suppress the DEB sensitivity of Hrq1-K318A by destabilizing the protein and mimicking *hrq1Δ* cells.

### Disruption of DNA binding leads to suppression of sensitivity to DEB

#Figure 2A and 2B show that the *hrq1-K318A* second-site suppressors that we collected largely phenocopy the DEB sensitivity level of *hrq1Δ* cells. Conceptually, it is simple for a mutant protein to behave like a null if the mutation destabilizes the helicase, and it is quickly degraded. Our western blotting (Fig. 2C and 2D) and computational analyses (Fig. 3A and Table S3) indicate that this may be a common route of DEB sensitivity suppression, with the Hrq1-K318A,C979Y and -K318A,A1006P suppressor mutants being characterized examples.

Unlike the K318A,C979Y and K318A,A1006P suppressors, the Hrq1-K318A,P732S mutant is expressed at approximately wild-type levels *in vivo* (Fig. 2C and 2D), arguing against protein instability as the only route to the suppression of DEB sensitivity. However, if the suppressor mutation ablated the helicase’s nuclear localization, this could also phenocopy *hrq1Δ* rather than *hrq1-K318A*. The nuclear localization sequence (NLS) of Hrq1 has not been identified, but cNLS Mapper (76) predicts that it exists from amino acids 932-942 (ERQTKRKRPAR). This motif is not impacted by any of the suppressor mutations that we isolated, other than those with frameshifts and/or early stop codons (Table 2). However, even if the predicted NLS is wrong, lack of nuclear localization would still be an uncommon route of suppressor generation because suppressor mutations were isolated throughout the length of the protein (Fig. 2E) rather than near a single putative NLS motif.

Therefore, we next hypothesized that mutations that impact the DNA binding of Hrq1-K318A could function as another way to yield suppressors. To test this hypothesis, we attempted to generate recombinant Hrq1-K318,P732S, -K318A,C979Y, and -K318A,A1006P mutant proteins to analyse their ability to bind ssDNA. Unfortunately, despite repeated attempts, we failed to produce useful amounts of the K318A,C979Y and K318A,A1006P proteins. This is likely due to their apparent instability in yeast (Fig. 2C and 2D). However, we did succeed in purifying Hrq1-P732S and Hrq1-K318A,P732S proteins (Fig. 4A).

**Figure 4.**
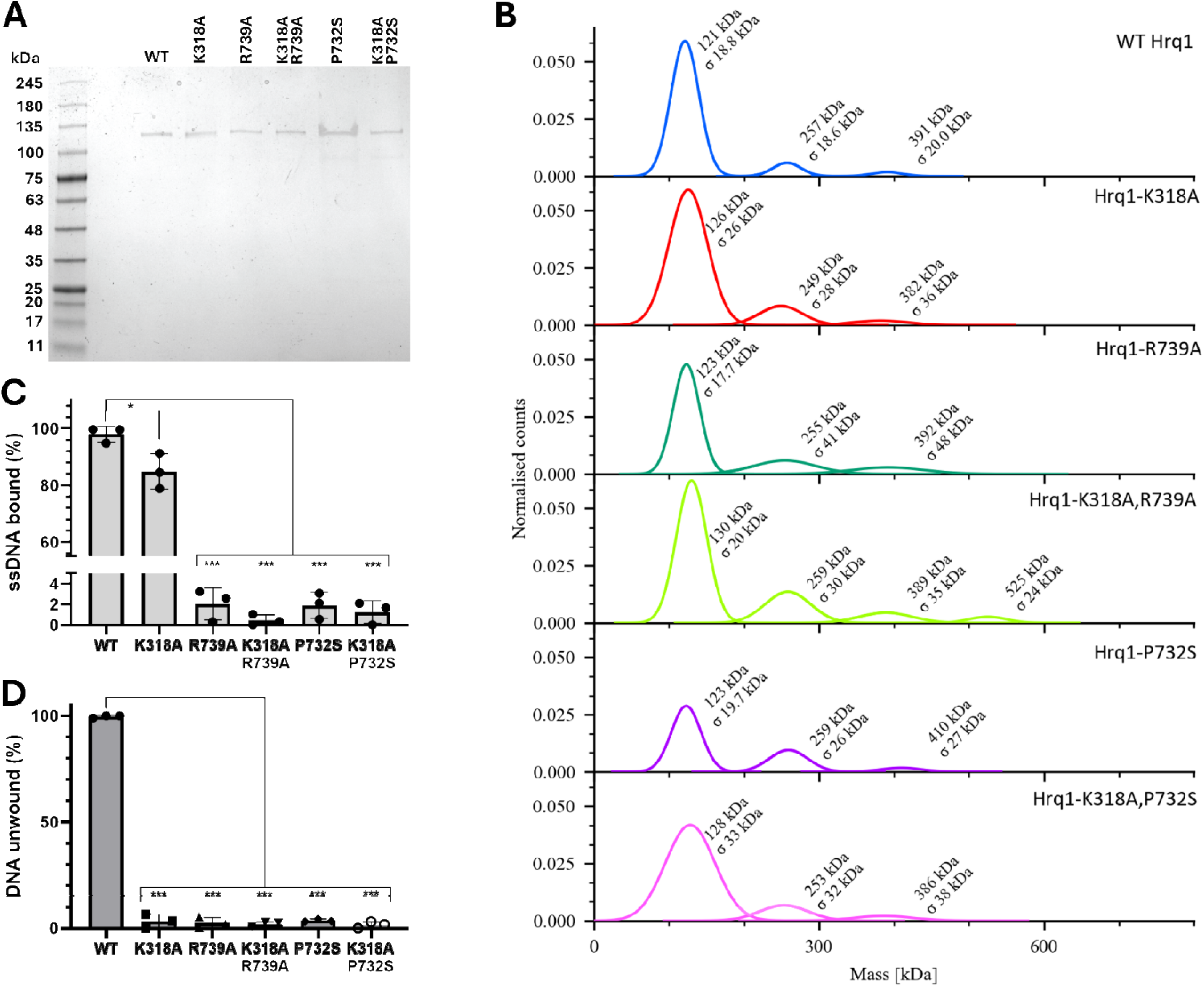
Ablating Hrq1-K318A DNA binding leads to suppression of ICL sensitivity. **A)** Recombinant wild-type (WT) and single or double-mutant Hrq1 protein preparations. Coomassie-stained SDS-PAGE gel of 2 ng of each of the indicated proteins run alongside a molecular weight marker ladder. **B)** Recombinant wild-type and mutant Hrq1 preparations form multimers in solution. Mass ph tometry profiles indicate that purified Hrq1 and mutant proteins (∼130 kDa) form dimers (∼260 kDa), trimers (∼390 kDa), and tetramers (∼520 kDa). The average molecular weight of single molecules in each Gaussian curve is listed, as well as the standard deviation (σ). **C)** DNA binding activity and **D)** helicase activity of Hrq1 and mutant proteins. Gel shift assays (n = 3) using 200 nM protein and 2 nM ssDNA probe were performed at 30°C to analyse ssDNA binding. Helicase assays used a fork substrate and included ATP to catalyse translocation and unwinding. In both c and D, the individual values are shown for each protein, as well as the average (bars) and standard deviation (error bars). One-way ANOVAs were used to determine significance. *, *p* < 0.05; and ***, *p* < 0.001.

To ensure that these mutants were properly folded and did not form soluble micro-aggregates, we analysed them by mass photometry (Fig. 4B). WT recombinant Hrq1 is known to oligomerize, with the proportion of different oligomers dependent upon the expression host (25,30). Here, mass photometry revealed the existence of monomers, dimers, trimers, and in some cases, tetramers for our WT and mutant preparations (Fig. 4B). As typical with baculovirus expression of Hrq1 in insect cells (30), the monomeric form dominated. Importantly, recombinant Hrq1-P732S and Hrq1-K318A,P732S displayed the same mass photometry profiles as WT and Hrq1-K318A (Fig. 4B). When these proteins are denatured with heat and SDS, the oligomeric peaks disappear, and only monomeric protein is evident (*e.g.*, Fig. S8).

Having demonstrated that our Hrq1-P732S and Hrq1-K318A,P732S preparations are stably folded, we next assessed their ability to bind ssDNA. We previously reported that Hrq1-K318A can bind to ssDNA but with approximately twofold lower affinity than Hrq1 (32), and that result is recapitulated here with an IR-labelled poly(dT) 30mer ssDNA (*p* = 0.0015) (Fig. 4C). In comparison, ssDNA binding by recombinant Hrq1-P732S and Hrq1-K318A,P732S was markedly reduced compared to WT Hrq1 and Hrq1-K318A (*p* < 0.0001). To further characterize these mutant proteins, we also tested their ability to unwind DNA. Unsurprisingly, for proteins that lacked ssDNA binding activity, Hrq1-P732S and Hrq1-K318A,P732S also lacked helicase activity (Fig. 4D). These data are consistent with the hypothesis that knocking out the DNA binding of Hrq1-K318A should relieve its dominant negative activity and phenocopy an Hrq1 null.

### *De novo* design of a suppressor

As another test of the hypothesis that disrupting the DNA binding activity of Hrq1-K318A is a route of DEB sensitivity suppression, we attempted to design a DNA binding mutant of Hrq1/Hrq1-K318A *de novo*. Assuming that the hypothesis is correct, cells expressing the double mutant with K318A should be resistant to DEB-induced ICL damage. We recently reported that an N-terminal truncation mutant of Hrq1 (Hrq1ΔN) largely lacks ssDNA binding activity (36). Therefore, we combined the ΔN allele with K318A (full-length Hrq1 residue numbering) to create a yeast strain expressing a Hrq1ΔN-K318A mutant and tested it for ICL sensitivity. We found that *hrq1ΔN-K318A* cells are less sensitive to DEB than the *hrq1-K318A* strain, growing more like *hrq1Δ* cells (Fig. S9). Thus, to a first approximation, inhibiting DNA binding in the Hrq1-K318A mutant does suppress DEB sensitivity. However, the comparison of a full-length Hrq1 mutant to this truncation is problematic, especially when considering the Hrq1-Pso2 ICL repair pathway and the putative involvement of the N-termini of both enzymes (29).

To determine if we could rationally design a full-length *hrq1-K318A* second-site suppressor, we turned back to the MrfA enzyme. Several residues in the *B. subtilis* helicase are demonstrated to be involved in ssDNA binding (62), many of which are conserved in *S. cerevisiae* Hrq1. This includes MrfA R491 (R739 in Hrq1; Fig. 5A), which decreases ssDNA binding by MrfA when mutated to an alanine 62). Based on homology, we hypothesized that the R739A mutation in Hrq1 should likewise reduce ssDNA binding. We generated recombinant Hrq1-R739A and Hrq1-K318A,R739A proteins (Fig. 4A) and tested their ssDNA binding activity using gel shifts. Compared to WT Hrq1 and Hrq1-K318A, the ability to bind ssDNA by Hrq1-R739A was severely decreased (Fig. 4C and 5B). Similarly, the recombinant Hrq1-K318A, R739A double mutant also lacked DNA binding activity (Fig. 4C).

**Figure 5.**
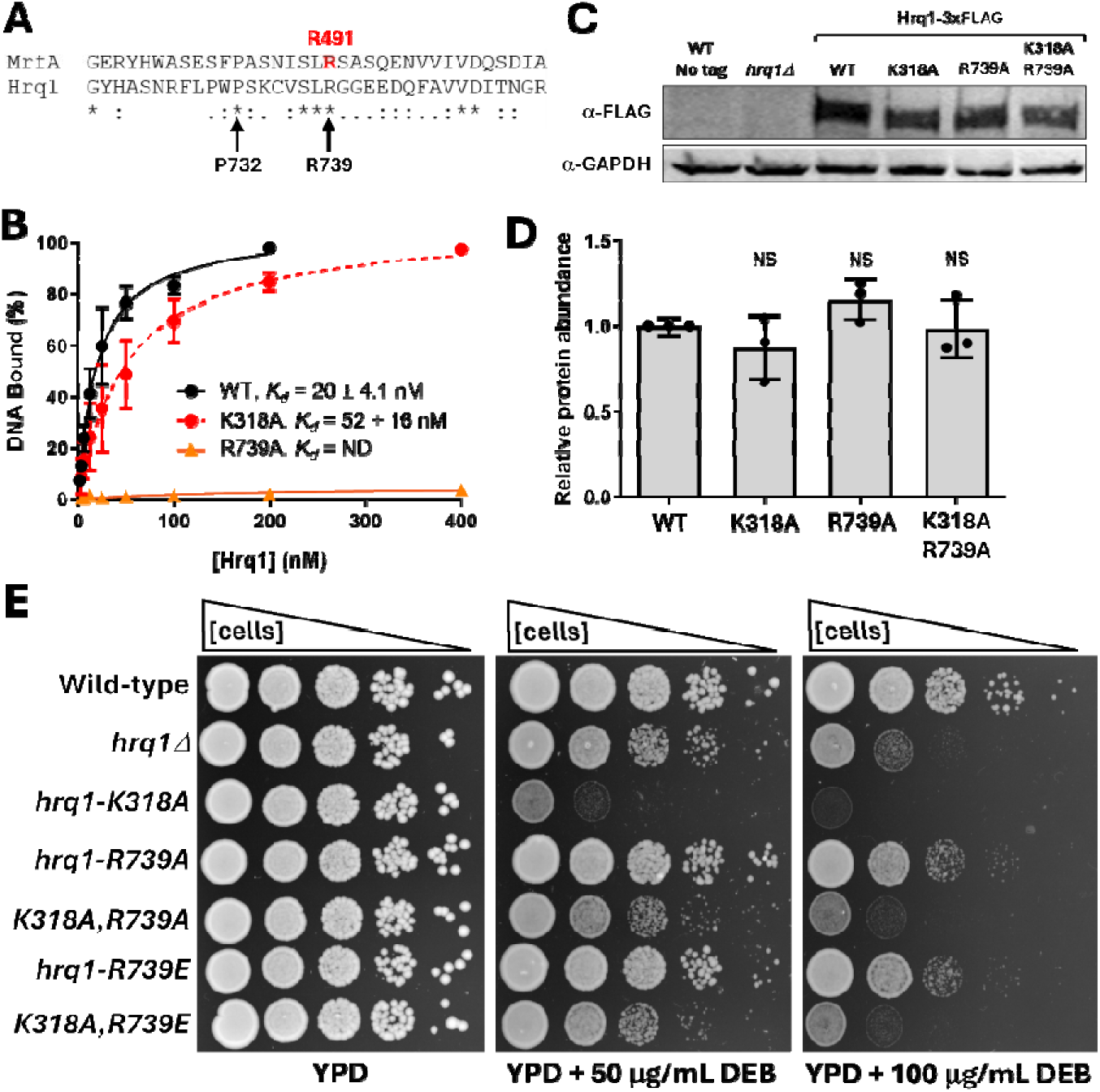
The R739A mutation disrupts Hrq1 ssDNA binding. **A)** Protein alignment of a conserved peptide in *B. subtilis* MrfA and *S. cerevisiae* Hrq1. MrfA R491 is noted in red, and the Hrq1 P732 and R739 residues are shown. Identical (*), strongly conserved (:), and conserved (.) residues are marked. **B)** Neutralizing R739 negatively impacts Hrq1 ssDNA binding. Gel shift as ays (n = 3) using 0-200 or 0-400 nM protein and 2 nM ssDNA probe were performed for the three indicated proteins. Binding was quantified, and the averages and standard deviations are plotted. Binding isotherms were fitted to each dataset to calculate the binding affinity (K_d_), which could not be determined (ND) f r Hrq1-R739A due to its poor binding. **C)** Western blot of 3xFLAG-tagged WT and mutant protein levels in yeast. GAPDH was used as a loading control. **D)** The amounts of mutant Hrq1 proteins in yeast did significantly differ compared to WT in n = 3 western blots as determined by one-way ANOVA. NS, *p* > 0.01 **E)** The *hrq1-K318A,R739A* and *hrq1-K318A,R739E* mutants are suppressors of DEB sensitivity.

Having proven that the R739A mutation negatively impacts ssDNA binding, we next introduced this mutation *in vivo*. Both the Hrq1-R739A and Hrq1-K318A,R739A mutants are produced at WT levels in yeast (Fig. 5D), so we also tested them for DEB sensitivity. As shown in Figure 5E, the *hrq1-K318A,R739A* double mutant is more resistant to DEB damage than *hrq1-K318A*, again confirming that disrupting ssDNA binding by the K318A mutant suppresses sensitivity to DEB. The *hrq1-R739A* single mutant was slightly less sensitive to DEB than the *hrq1Δ* allele, suggesting that perhaps Hrq1-R739A retains some level of ssDNA binding *in vivo*. In an attempt to make a stronger DNA binding-defective allele, we also created *hrq1-R739E* and *hrq1-K318A,R739E* strains, reasoning that the charge swap may repel the DNA. As above, the double mutant is a DEB sensitivity suppressor, and the single mutant grows better than *hrq1Δ* on DEB plates (Fig 5E). Because DNA binding-impaired alleles of *HRQ1* suppress the dominant negative phenotype of *hrq1-K318A* but do not fully phenocopy *hrq1Δ* in DEB sensitivity assays, these data suggest that the Hrq1-K318A protein acts as a dominant negative not just due to non-productive DNA binding. This phenomenon and others are discussed below.

## DISCUSSION

DEB exposure causes ICLs, which pose a severe threat to genome stability by covalently linking the two strands of DNA and blocking replication and transcription (40). In *S. cerevisiae*, repair of these lesions relies heavily on the Hrq1 helicase and the Pso2 nuclease, with Hrq1 stimulating Pso2-mediated processing of ICLs (25,28–30,40). In this study, we combined genetic suppressor analysis, computational modelling, and biochemical characterization to investigate the consequences of disabling Hrq1 helicase activity. Our results demonstrate that the dominant-negative phenotype of *hrq1-K318A* can be suppressed by intragenic mutations that either reduce protein abundance or impair DNA binding.

### Second-site suppressors in Hrq1-K318A

The spontaneous suppressor screen yielded an unexpected but highly informative result: suppression of *hrq1-K318A* sensitivity to DEB-induced ICL damage occurred overwhelmingly through secondary mutations within the *hrq1-K318A* gene itself. Both whole-genome sequencing and targeted amplicon sequencing demonstrated that the majority of suppressors carried second-site mutations distributed throughout the Hrq1 coding sequence (Fig. 2 and Table 2). Reversion of the K318A mutation would be expected to suppress DEB sensitivity, but all suppressor clones retained the K318A substitution in addition to one or more additional alterations, indicating that suppression was not due to restoration of catalytic activity. Reconstruction of representative mutations confirmed that suppression is monogenic and directly attributable to altered Hrq1 function (Fig. 2B), rather than to the cumulative effects of multiple DEB-induced mutations elsewhere in the genome.

Consistent with previous observations (25,29,30,36), *hrq1-K318A* cells were more sensitive to DEB than *hrq1*Δ cells (Fig. 2, 3, 5, S1, S2, and S9), supporting the conclusion that *hrq1-K318A* functions as a dominant-negative allele. This interpretation is further supported by complementation experiments showing that expression of WT *HRQ1* efficiently rescued the DEB sensitivity of *hrq1*Δ cells but only weakly alleviated the sensitivity of *hrq1-K318A* cells (Fig. S4). Together, these findings support a model in which Hrq1-K318A is recruited to sites of ICL damage but, lacking ATPase and helicase activity, interferes with or requires a less efficient, alternative repair mechanism that would otherwise partially compensate for complete loss of Hrq1–Pso2 function (Fig. 6).

**Figure 6.**
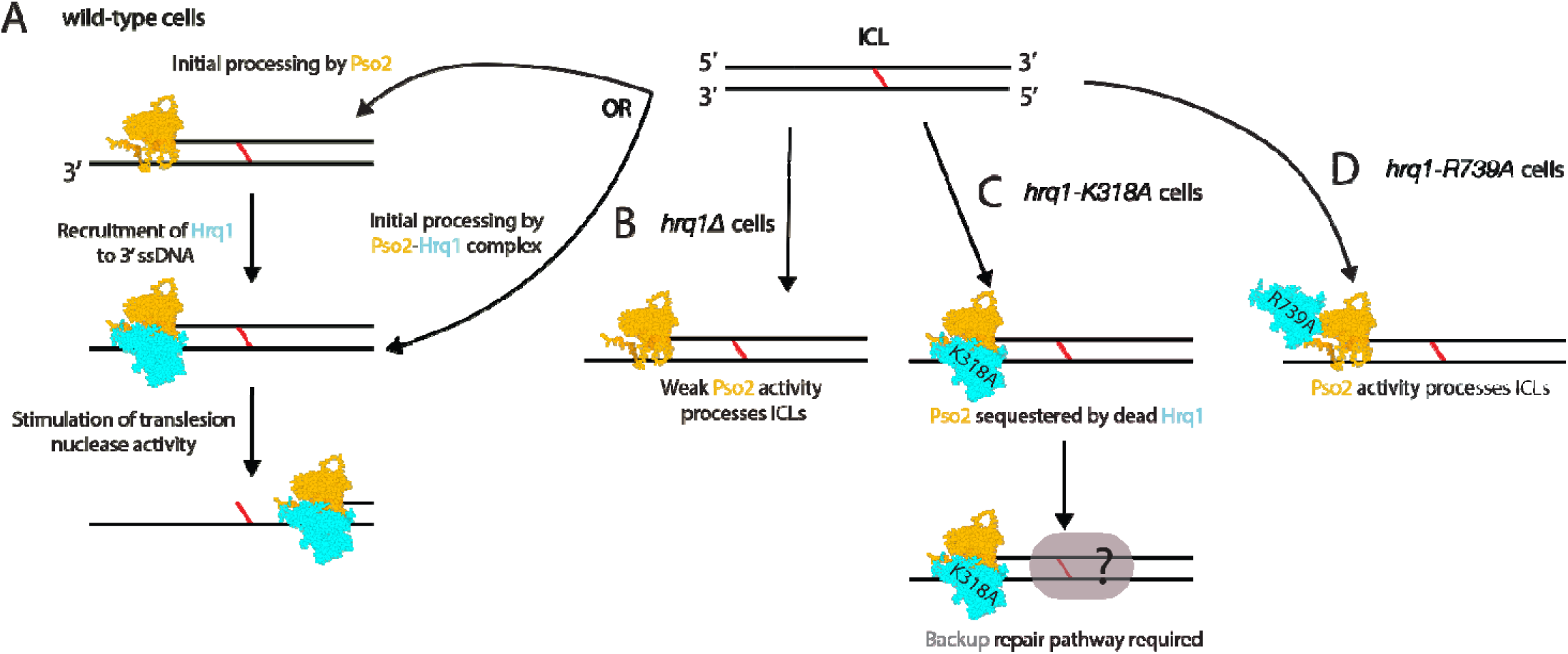
Working model for ICL repair in wild-type and *hrq1* mutant cells. **A)** When an ICL lesion is formed, Pso2 may initially recognize it or be recruited to it alone to begin degrading one strand of DNA from a 5⍰ phosphate (left). This would generate a ssDNA substrate for Hrq1 to bind to for 3⍰-5⍰ translocation. Alternatively, Hrq1 and Pso2 may be recruited as a single unit for initial processing (right). Both paths lead to stimulation of Pso2 nuclease activity by Hrq1 for ssDNA digestion through the lesion. B) In *hrq1Δ* cells, Pso2 is still recruited to repair the ICL but does so weakly in the absence of Hrq1 stimulation. C) In *hrq1-K318A* cells, Pso2 association with the non-translocating Hrq1 may interfere with Pso2-mediated repair, necessitating compensatory repair mechanisms. Thus, *hrq1-K318A* cells are more sensitive to ICL damage than *hrq1Δ* cells because they largely phenocopy *pso2Δ*. D) In *hrq1-R739A* cells or other mutants in which Hrq1 lacks ssDNA binding activity, Hrq1 still binds to Pso2, but Pso2 is not locked down on the DNA and can still process ICLs. This model summarizes interpretations of genetic and biochemical data but does not represent a directly tested mechanism.

### Mutational modulation of a catalytically inactive Hrq1 scaffold

The recovery of Hrq1 variants that suppress DEB sensitivity in the *hrq1-K318A* background suggests that Hrq1 possesses critical non-catalytic roles in the response to DEB damage. Because the K318A mutation abolishes ATP hydrolysis and DNA translocation, the suppression observed here cannot be attributed to the restoration or tuning of helicase motor activity. Instead, these suppressors likely modulate the physical properties of the Hrq1 scaffold itself, potentially altering its stability, how the inactive protein occupies DNA lesions, or how it interacts with the broader repair machinery.

Our computational analyses reveal a notable divergence between evolutionary conservation and structural stability within the catalytic core (helicase and RQC domains). Residues such as R442, V641, and H683 are identified by ESM-1v as being under intense purifying selection, yet their substitution is predicted to be structurally neutral by MAESTRO (Fig. 3A and Table S3). In a WT enzyme, these residues may be important for catalysis. However, in the Hrq1-K318A “dead-motor” context, we propose that these conserved residues act as high-affinity tethers that cause the inactive helicase to remain stably – but unproductively – bound to DEB ICL repair intermediates.

By substituting these evolutionarily conserved residues without triggering global misfolding, as indicated by the marginal ΔΔG values, these suppressor mutations may decrease the stability of the Hrq1-DNA complex. This “loosening” of the inactive scaffold could prevent Hrq1-K318A from acting as a physical roadblock, thereby facilitating the recruitment of auxiliary helicases or nucleases that can process the DEB-induced damage in an Hrq1-independent manner.

A second cluster of suppressors localized near the RHCD region exhibited higher consensus across all models, being flagged as both evolutionarily deleterious and structurally destabilizing (Fig. 3A and Table S3). Unlike the catalytic core, the RHCD appears to require strict structural integrity to maintain its function. Given the lack of motor activity in the *hrq1-K318A* background, the RHCD may globally impact protein stability or serve as a critical docking surface for regulatory factors. The destabilization of this domain via mutations like C979Y or A1006P might reconfigure the Hrq1 interactome (34), potentially releasing stalled repair complexes or altering the signalling hierarchy at DEB lesions.

Collectively, these findings suggest that Hrq1’s contribution to genomic stability during DEB repair is bifurcated: while its helicase activity is normally paramount, the physical occupancy and scaffolding properties of the protein are equally vital for ensuring that DNA repair intermediates are efficiently processed and/or handed off to downstream pathways. The lack of consensus among the computational methods used to characterize our suppressors suggests that some of them are mechanistically surgical. They are not all simply destabilizing the protein (which MAESTRO would have flagged); they are specifically tuning evolutionary conserved residues (Table 2) required for DNA processing.

### Disruption of Hrq1 ssDNA binding

An apparent discrepancy in our model of Hrq1-K318A as a dominant negative arises from the behaviour of the mutants that impair ssDNA binding. Recombinant Hrq1-P732S and Hrq1-R739A proteins display markedly reduced ssDNA binding and helicase activity *in vitro* compared to WT Hrq1 and Hrq1-K318A (Fig. 4C-D and 5B). Likewise, introducing either mutation into the *hrq1-K318A* background suppresses sensitivity to DEB-induced ICL damage (Fig. 3B, 5E, and S2), consistent with the idea that DNA engagement is required for Hrq1-K318A dominance.

However, despite their strong biochemical defects *in vitro*, the *hrq1-P732S*, *hrq1-R739A*, and *hrq1-R739E* single-mutant strains do not phenocopy *hrq1*Δ *in vivo*. Instead, these alleles display an intermediate DEB sensitivity between *hrq1*Δ and *hrq1-K318A* (Fig. 3B and 5E). This observation suggests that the *in vitro* ssDNA binding and helicase assays do not fully capture the functional requirements for Hrq1 activity during ICL repair in cells.

One explanation is that Hrq1 operates within multiprotein repair assemblies , including complexes containing the Pso2 nuclease (29,30). In this context, protein-protein interactions and cooperative engagement of repair intermediates may stabilize transient DNA association even when intrinsic Hrq1 ssDNA binding affinity is weakened. Additionally, the short homopolymeric ssDNA substrates used may not accurately reflect the DNA structures encountered during ICL repair, such as partially processed lesions, forked intermediates, or protein-bound DNA substrates. Thus, mutations such as P732S and R739A may preferentially weaken nonproductive DNA engagement while still permitting sufficient repair-associated interactions to support partial Hrq1 function *in vivo*.

These findings suggest that dominant-negative toxicity depends more strongly on stable, nonproductive DNA engagement by Hrq1-K318A than on complete retention of Hrq1 biochemical activity. More broadly, they highlight an important distinction between DNA binding measured under defined biochemical conditions and the functional requirements for Hrq1-dependent ICL repair in cells.

### Working model for ICL repair in yeast

Collectively, these findings support a model in which DNA binding without catalytic activity converts Hrq1 into a toxic roadblock during ICL repair (Fig. 6). In this model, Hrq1-K318A is recruited to ICL-associated DNA structures but cannot translocate or remodel the substrate, thereby blocking access by compensatory repair enzymes or requiring non-optimal backup repair mechanisms (Fig. 6B). Secondary mutations that destabilize Hrq1 or prevent its association with DNA alleviate this blockade, allowing alternative repair pathways to function and restoring resistance to ICL damage (Fig. 6C). This framework explains why *hrq1-K318A* mutants are more deleterious than *hrq1*Δ mutants and why suppression can be achieved through a wide variety of intragenic second-site mutations.

Beyond RecQ4-family helicases, our findings may reflect a broader principle in genome maintenance pathways: catalytically impaired helicases can remain biologically active through persistent nucleic acid or protein interactions, thereby interfering with repair or replication processes in a dominant-negative manner. Similar concepts have emerged in other helicase systems in which catalytic activity is partially separable from structural or regulatory functions. For example, the nuclease-helicase Dna2 retains essential genome maintenance functions even when helicase activity is compromised (77,78), while the Fe-S helicase RTEL1 additionally relies on specialized recruitment and protein interaction functions that are mechanistically distinct from helicase translocation itself (79,80). Likewise, the DExH-box protein DHX58/LGP2 functions primarily as an RNA-binding regulator of antiviral signalling rather than as a processive helicase enzyme (81–83). Together, these observations support the idea that incomplete loss-of-function helicase alleles may perturb cellular pathways through aberrant substrate engagement or scaffolding interactions rather than through simple loss of catalytic activity alone.

Although our study is focused on *S. cerevisiae* Hrq1, the mechanistic principles described here are likely relevant to RecQ4-family helicases more broadly. Human RECQL4 plays essential roles in genome maintenance, and pathogenic RECQ4 variants may retain partial expression or DNA binding capacity while exhibiting reduced catalytic activity (18,21,23,26,38,84–90). Our findings suggest that such alleles may be deleterious not simply because of loss of helicase function or destabilization of the mutant protein, but because DNA engagement by catalytically impaired proteins can actively interfere with repair processes. By separating the contributions of DNA binding and enzymatic activity, this work provides a conceptual framework for understanding how incomplete loss-of-function mutations in RecQ4-family helicases may have more severe consequences than null alleles. Indeed, many of the Hrq1 suppressor mutations targeted residues that are conserved in RECQL4 and are themselves mutated in tumour samples (72). This suggests that using small molecules to target DNA binding by catalytically inactive disease alleles of human RECQL4 is a viable treatment strategy to mitigate the pathogenic effects if the helicase mutation.

In summary, this study reveals DNA binding as the central determinant of dominant-negative Hrq1 activity during ICL repair and demonstrates that disrupting this interaction suppresses repair defects caused by catalytic inactivation. These findings refine our understanding of Hrq1 function, illuminate the hazards of helicase engagement without catalysis, and provide mechanistic insight into how RecQ4-family helicases contribute to genome stability.

## DATA AVAILABILITY

The data underlying this article are available in the article, in its online supplementary material, or will be shared upon reasonable request to the corresponding author.

## SUPPLEMENTARY DATA

Supplementary Data are available at NAR online.

## ACKNOWLEDGEMENTS

We thank members of the Bochman lab for providing insightful feedback during the course of this work. We wish to acknowledge and honour the myaamiaki, Lënape, Bodwéwadmik, and saawanwa people, on whose ancestral homelands and resources Indiana University Bloomington is built.

## AUTHOR CONTRIBUTIONS

Robert Simmons: Conceptualization, Formal analysis, Methodology, Validation, Writing—original draft. Faith E. McDevitt: Formal analysis, Methodology, Validation, Writing— review & editing. Alexandra Hurlock: Formal analysis, Methodology, Validation, Writing— review & editing. Michael E. Kumcu: Formal analysis, Methodology, Validation, Writing— review & editing. Matthew Bochman: Conceptualization, Formal analysis, Methodology, Validation, Writing— review & editing.

## REFERENCES

1. Croteau, D.L., Popuri, V., Opresko, P.L. and Bohr, V.A. (2014) Human RecQ helicases in DNA repair, recombination, and replication. Annu Rev Biochem, 83, 519–552.

2. Abu-Libdeh, B., Jhujh, S.S., Dhar, S., Sommers, J.A., Datta, A., Longo, G.M., Grange, L.J., Reynolds, J.J., Cooke, S.L., McNee, G.S. et al. (2022) RECON syndrome is a genome instability disorder caused by mutations in the DNA helicase RECQL1. J Clin Invest, 132.

3. Larizza, L., Roversi, G. and Volpi, L. (2010) Rothmund-Thomson syndrome. Orphanet J Rare Dis, 5, 2.

4. Lee, J.W., Harrigan, J., Opresko, P.L. and Bohr, V.A. (2005) Pathways and functions of the Werner syndrome protein. Mech. Ageing Dev., 126, 79–86.

5. Langer, K., Cunniff, C.M. and Kucine, N. (1993) In Adam, M. P., Bick, S., Mirzaa, G. M., Pagon, R. A., Wallace, S. E. and Amemiya, A. (eds.), GeneReviews((R)), Seattle (WA).

6. Mojumdar, A. (2020) Mutations in conserved functional domains of human RecQ helicases are associated with diseases and cancer: A review. Biophys Chem, 265, 106433.

7. Gupta, P., Majumdar, A.G. and Patro, B.S. (2022) Enigmatic role of WRN-RECQL helicase in DNA repair and its implications in cancer. J Transl Genet Genom, 6, 147–156.

8. Baltgalvis, K.A., Lamb, K.N., Symons, K.T., Wu, C.C., Hoffman, M.A., Snead, A.N., Song, X., Glaza, T., Kikuchi, S., Green, J.C. et al. (2024) Chemoproteomic discovery of a covalent allosteric inhibitor of WRN helicase. Nature, 629, 435–442.

9. Ferretti, S., Hamon, J., de Kanter, R., Scheufler, C., Andraos-Rey, R., Barbe, S., Bechter, E., Blank, J., Bordas, V., Dammassa, E., et al. (2024) Discovery of WRN inhibitor HRO761 with synthetic lethality in MSI cancers. Nature, 629, 443–449.

10. van Wietmarschen, N., Sridharan, S., Nathan, W.J., Tubbs, A., Chan, E.M., Callen, E., Wu, W., Belinky, F., Tripathi, V., Wong, N., et al. (2020) Repeat expansions confer WRN dependence in microsatellite-unstable cancers. Nature, 586, 292–298.

11. Behan, F.M., Iorio, F., Picco, G., Goncalves, E., Beaver, C.M., Migliardi, G., Santos, R., Rao, Y., Sassi, F., Pinnelli, M. et al. (2019) Prioritization of cancer therapeutic targets using CRISPR-Cas9 screens. Nature, 568, 511–516.

12. Chan, E.M., Shibue, T., McFarland, J.M., Gaeta, B., Ghandi, M., Dumont, N., Gonzalez, A., McPartlan, J.S., Li, T., Zhang, Y. et al. (2019) WRN helicase is a synthetic lethal target in microsatellite unstable cancers. Nature, 568, 551–556.

13. Chen, J., Situ, Y., Cai, J., Liao, W.Y., Jiang, D.B., Huang, C.S., Zheng, J.C., Peng, L.J. and Lin, H. (2026) Upregulated BLM and RECQL4 in Osteosarcoma: Association with Poor Prognosis, Immune Cell Infiltration, and Inhibitory Effects of Sphingosine Kinase 1 Inhibitor II/Pilaralisib. Immunotargets Ther, 15.

14. Ma, X., Tian, F., Xiao, Y., Huang, M., Song, D., Chen, X. and Xu, H. (2024) Synergistic effects of bloom helicase (BLM) inhibitor AO/854 with cisplatin in prostate cancer. Sci Rep, 14, 24962.

15. Ovejero, S., Viziteu, E., Dutrieux, L., Devin, J., Lin, Y.L., Alaterre, E., Jourdan, M., Basbous, J., Requirand, G., Robert, N. et al. (2022) The BLM helicase is a new therapeutic target in multiple myeloma involved in replication stress survival and drug resistance. Front Immunol, 13, 983181.

16. Ma, X.Y., Xu, H.Q., Zhao, J.F., Ruan, Y. and Chen, B. (2022) Discovery of a Novel Bloom’s Syndrome Protein (BLM) Inhibitor Suppressing Growth and Metastasis of Prostate Cancer. Int J Mol Sci, 23.

17. Kaczmarczyk, A., Sokolowski, M., Wojnicki, K., Pabis, M., Wojtas, B., Ciechomska, I.A., Poleszak, K., Gielniewski, B., Krol, S.K., Guille, M. et al. (2025) Novel mutations in the RECQL4 gene affect its helicase functions, interactions with the BLM helicase and chemotherapeutics-induced cell death. Cell Death Discov, 11, 560.

18. Balajee, A.S. (2021) Human RecQL4 as a Novel Molecular Target for Cancer Therapy. Cytogenet Genome Res, 161, 305–327.

19. Yokoyama, H., Moreno-Andres, D., Astrinidis, S.A., Hao, Y., Weberruss, M., Schellhaus, A.K., Lue, H., Haramoto, Y., Gruss, O.J. and Antonin, W. (2019) Chromosome alignment maintenance requires the MAP RECQL4, mutated in the Rothmund-Thomson syndrome. Life Sci Alliance, 2.

20. Singh, D.K., Popuri, V., Kulikowicz, T., Shevelev, I., Ghosh, A.K., Ramamoorthy, M., Rossi, M.L., Janscak, P., Croteau, D.L. and Bohr, V.A. (2012) The human RecQ helicases BLM and RECQL4 cooperate to preserve genome stability. Nucleic Acids Res, 40, 6632–6648.

21. Su, Y., Meador, J.A., Calaf, G.M., Proietti De-Santis, L., Zhao, Y., Bohr, V.A. and Balajee, A.S. (2010) Human RecQL4 helicase plays critical roles in prostate carcinogenesis. Cancer Res, 70, 9207–9217.

22. Kamimura, Y., Masumoto, H., Sugino, A. and Araki, H. (1998) Sld2, which interacts with Dpb11 in Saccharomyces cerevisiae, is required for chromosomal DNA replication. Mol Cell Biol, 18, 6102–6109.

23. Xu, X., Chang, C.W., Li, M., Liu, C. and Liu, Y. (2021) Molecular Mechanisms of the RECQ4 Pathogenic Mutations. Front Mol Biosci, 8, 791194.

24. Abe, T., Yoshimura, A., Hosono, Y., Tada, S., Seki, M. and Enomoto, T. (2011) The N-terminal region of RECQL4 lacking the helicase domain is both essential and sufficient for the viability of vertebrate cells. Role of the N-terminal region of RECQL4 in cells. Biochim Biophys Acta, 1813, 473–479.

25. Bochman, M.L., Paeschke, K., Chan, A. and Zakian, V.A. (2014) Hrq1, a homolog of the human RecQ4 helicase, acts catalytically and structurally to promote genome integrity. Cell Rep, 6, 346–356.

26. Macris, M.A., Krejci, L., Bussen, W., Shimamoto, A. and Sung, P. (2006) Biochemical characterization of the RECQ4 protein, mutated in Rothmund-Thomson syndrome. DNA Repair (Amst), 5, 172–180.

27. Barea, F., Tessaro, S. and Bonatto, D. (2008) In silico analyses of a new group of fungal and plant RecQ4-homologous proteins. Comput Biol Chem, 32, 349–358.

28. Rogers, C.M. and Bochman, M.L. (2017) Saccharomyces cerevisiae Hrq1 helicase activity is affected by the sequence but not the length of single-stranded DNA. Biochem Biophys Res Commun, 486, 1116–1121.

29. Rogers, C.M., Lee, C.Y., Parkins, S., Buehler, N.J., Wenzel, S., Martinez-Marquez, F., Takagi, Y., Myong, S. and Bochman, M.L. (2020) The yeast Hrq1 helicase stimulates Pso2 translesion nuclease activity and thereby promotes DNA interstrand crosslink repair. J Biol Chem, 295, 8945–8957.

30. Rogers, C.M., Wang, J.C., Noguchi, H., Imasaki, T., Takagi, Y. and Bochman, M.L. (2017) Yeast Hrq1 shares structural and functional homology with the disease-linked human RecQ4 helicase. Nucleic Acids Res, 45, 5217–5230.

31. Nickens, D.G. and Bochman, M.L. (2022) Genetic and biochemical interactions of yeast DNA helicases. Methods, 204, 234–240.

32. Nickens, D.G., Rogers, C.M. and Bochman, M.L. (2018) The Saccharomyces cerevisiae Hrq1 and Pif1 DNA helicases synergistically modulate telomerase activity in vitro. J Biol Chem, 293, 14481–14496.

33. Nickens, D.G., Sausen, C.W. and Bochman, M.L. (2019) The Biochemical Activities of the Saccharomyces cerevisiae Pif1 Helicase Are Regulated by Its N-Terminal Domain. Genes (Basel), 10.

34. Rogers, C.M., Sanders, E., Nguyen, P.A., Smith-Kinnaman, W., Mosley, A.L. and Bochman, M.L. (2020) The Genetic and Physical Interactomes of the Saccharomyces cerevisiae Hrq1 Helicase. G3 (Bethesda), 10, 4347–4357.

35. Sanders, E., Nguyen, P.A., Rogers, C.M. and Bochman, M.L. (2020) Comprehensive Synthetic Genetic Array Analysis of Alleles That Interact with Mutation of the Saccharomyces cerevisiae RecQ Helicases Hrq1 and Sgs1. G3 (Bethesda), 10, 4359–4368.

36. Shumaker, K.A., Kumcu, M.E., McDevitt, F.E., Rogers, C.M. and Bochman, M.L. (2025) A screen for synthetic genetic interactions with the Saccharomyces cerevisiae hrq1DeltaN allele. G3 (Bethesda).

37. Luong, T.T., Li, Z., Priedigkeit, N., Parker, P.S., Bohm, S., Rapchak, K., Lee, A.V. and Bernstein, K.A. (2022) Hrq1/RECQL4 regulation is critical for preventing aberrant recombination during DNA intrastrand crosslink repair and is upregulated in breast cancer. PLoS Genet, 18, e1010122.

38. Jin, W., Liu, H., Zhang, Y., Otta, S.K., Plon, S.E. and Wang, L.L. (2008) Sensitivity of RECQL4-deficient fibroblasts from Rothmund-Thomson syndrome patients to genotoxic agents. Hum Genet, 123, 643–653.

39. Mo, D., Fang, H., Niu, K., Liu, J., Wu, M., Li, S., Zhu, T., Aleskandarany, M.A., Arora, A., Lobo, D.N. et al. (2016) Human Helicase RECQL4 Drives Cisplatin Resistance in Gastric Cancer by Activating an AKT-YB1-MDR1 Signaling Pathway. Cancer Res, 76, 3057–3066.

40. Rogers, C.M., Simmons III, R.H., Fluhler Thornburg, G.E., Buehler, N.J. and Bochman, M.L. (2020) Fanconi anemia-independent DNA inter-strand crosslink repair in eukaryotes. Prog Biophys Mol Biol.

41. Herskowitz, I. (1987) Functional inactivation of genes by dominant negative mutations. Nature, 329, 219–222.

42. Veitia, R.A. (2007) Exploring the molecular etiology of dominant-negative mutations. Plant Cell, 19, 3843–3851.

43. Meeks-Wagner, D. and Hartwell, L.H. (1986) Normal stoichiometry of histone dimer sets is necessary for high fidelity of mitotic chromosome transmission. Cell, 44, 43–52.

44. Lagna, G. and Hemmati-Brivanlou, A. (1998) Use of dominant negative constructs to modulate gene expression. Curr Top Dev Biol, 36, 75–98.

45. Stark, J.M., Hu, P., Pierce, A.J., Moynahan, M.E., Ellis, N. and Jasin, M. (2002) ATP hydrolysis by mammalian RAD51 has a key role during homology-directed DNA repair. J Biol Chem, 277, 20185–20194.

46. Pellegrini, L., Yu, D.S., Lo, T., Anand, S., Lee, M., Blundell, T.L. and Venkitaraman, A.R. (2002) Insights into DNA recombination from the structure of a RAD51-BRCA2 complex. Nature, 420, 287–293.

47. Wang, A.T., Kim, T., Wagner, J.E., Conti, B.A., Lach, F.P., Huang, A.L., Molina, H., Sanborn, E.M., Zierhut, H., Cornes, B.K. et al. (2015) A Dominant Mutation in Human RAD51 Reveals Its Function in DNA Interstrand Crosslink Repair Independent of Homologous Recombination. Mol Cell, 59, 478–490.

48. So, A., Dardillac, E., Muhammad, A., Chailleux, C., Sesma-Sanz, L., Ragu, S., Le Cam, E., Canitrot, Y., Masson, J.Y., Dupaigne, P., et al. (2022) RAD51 protects against nonconservative DNA double-strand break repair through a nonenzymatic function. Nucleic Acids Res, 50, 2651–2666.

49. George, J.W., Brosh, R.M., Jr. and Matson, S.W. (1994) A dominant negative allele of the Escherichia coli uvrD gene encoding DNA helicase II. A biochemical and genetic characterization. J Mol Biol, 235, 424–435.

50. Taneja, P., Gu, J., Peng, R., Carrick, R., Uchiumi, F., Ott, R.D., Gustafson, E., Podust, V.N. and Fanning, E. (2002) A dominant-negative mutant of human DNA helicase B blocks the onset of chromosomal DNA replication. J Biol Chem, 277, 40853–40861.

51. Wu, Y. and Brosh, R.M., Jr. (2010) Helicase-inactivating mutations as a basis for dominant negative phenotypes. Cell Cycle, 9, 4080–4090.

52. Sikorski, R.S. and Hieter, P. (1989) A system of shuttle vectors and yeast host strains designed for efficient manipulation of DNA in *Saccharomyces cerevisiae*. Genetics, 122, 19–27.

53. Longtine, M.S., McKenzie, A., 3rd, Demarini, D.J., Shah, N.G., Wach, A., Brachat, A., Philippsen, P. and Pringle, J.R. (1998) Additional modules for versatile and economical PCR-based gene deletion and modification in *Saccharomyces cerevisiae*. Yeast, 14, 953–961.

54. Larkin, M.A., Blackshields, G., Brown, N.P., Chenna, R., McGettigan, P.A., McWilliam, H., Valentin, F., Wallace, I.M., Wilm, A., Lopez, R. et al. (2007) Clustal W and Clustal X version 2.0. Bioinformatics, 23, 2947–2948.

55. von der Haar, T. (2007) Optimized protein extraction for quantitative proteomics of yeasts. PLoS One, 2, e1078.

56. Meier, J., Rao, R., Verkuil, R., Liu, J., Sercu, T. and Rives, A. (2021) Language models enable zero-shot prediction of the effects of mutations on protein function. Adv Neur In, 34.

57. Fleming, J., Magana, P., Nair, S., Tsenkov, M., Bertoni, D., Pidruchna, I., Lima Afonso, M.Q., Midlik, A., Paramval, U., Zidek, A. et al. (2025) AlphaFold Protein Structure Database and 3D-Beacons: New Data and Capabilities. J Mol Biol, 437, 168967.

58. Schymkowitz, J., Borg, J., Stricher, F., Nys, R., Rousseau, F. and Serrano, L. (2005) The FoldX web server: an online force field. Nucleic Acids Res, 33, W382–388.

59. Laimer, J., Hofer, H., Fritz, M., Wegenkittl, S. and Lackner, P. (2015) MAESTRO--multi agent stability prediction upon point mutations. BMC Bioinformatics, 16, 116.

60. Sievers, F., Wilm, A., Dineen, D., Gibson, T.J., Karplus, K., Li, W., Lopez, R., McWilliam, H., Remmert, M., Soding, J. et al. (2011) Fast, scalable generation of high-quality protein multiple sequence alignments using Clustal Omega. Mol Syst Biol, 7, 539.

61. Rice, P., Longden, I. and Bleasby, A. (2000) EMBOSS: the European Molecular Biology Open Software Suite. Trends Genet, 16, 276–277.

62. Roske, J.J., Liu, S., Loll, B., Neu, U. and Wahl, M.C. (2021) A skipping rope translocation mechanism in a widespread family of DNA repair helicases. Nucleic Acids Res, 49, 504–518.

63. Neuhoff, V., Arold, N., Taube, D. and Ehrhardt, W. (1988) Improved staining of proteins in polyacrylamide gels including isoelectric focusing gels with clear background at nanogram sensitivity using Coomassie Brilliant Blue G-250 and R-250. Electrophoresis, 9, 255–262.

64. Kang, D.H., Gho, Y.S., Suh, M.K. and Kang, C.H. (2002) Highly sensitive and fast protein detection with coomassie brilliant blue in sodium dodecyl sulfate-polyacrylamide gel electrophoresis. B Korean Chem Soc, 23, 1511–1512.

65. Nickens, D.G., Feng, Z., Shen, J., Gray, S.J., Simmons, R.H., 3rd, Niu, H. and Bochman, M.L. (2024) Cdc13 exhibits dynamic DNA strand exchange in the presence of telomeric DNA. Nucleic Acids Res, 52, 6317–6332.

66. Nickens, D.G., Gray, S.J., Simmons, R.H., 3rd and Bochman, M.L. (2025) Dimerization of Cdc13 is essential for dynamic DNA exchange on telomeric DNA. J Biol Chem, 301, 110496.

67. Simmons, R.H., 3rd, Rogers, C.M. and Bochman, M.L. (2021) A deep dive into the RecQ interactome: something old and something new. Curr Genet, 67, 761–767.

68. Fusco, D., Gralka, M., Kayser, J., Anderson, A. and Hallatschek, O. (2016) Excess of mutational jackpot events in expanding populations revealed by spatial Luria-Delbruck experiments. Nat Commun, 7, 12760.

69. Ellison, M.A., Walker, J.L., Ropp, P.J., Durrant, J.D. and Arndt, K.M. (2020) MutantHuntWGS: A Pipeline for Identifying Saccharomyces cerevisiae Mutations. G3 (Bethesda), 10, 3009–3014.

70. Bochman, M.L. (2014) Roles of DNA helicases in the maintenance of genome integrity. Mol Cell Oncol, 1, e963429.

71. Burby, P.E. and Simmons, L.A. (2019) A bacterial DNA repair pathway specific to a natural antibiotic. Mol Microbiol, 111, 338–353.

72. Tate, J.G., Bamford, S., Jubb, H.C., Sondka, Z., Beare, D.M., Bindal, N., Boutselakis, H., Cole, C.G., Creatore, C., Dawson, E. et al. (2019) COSMIC: the Catalogue Of Somatic Mutations In Cancer. Nucleic Acids Res, 47, D941–D947.

73. Van Maldergem, L., Piard, J., Larizza, L. and Wang, L.L. (1993) In Adam, M. P., Feldman, J., Mirzaa, G. M., Pagon, R. A., Wallace, S. E. and Amemiya, A. (eds.), GeneReviews((R)), Seattle (WA).

74. Vargas, F.R., de Almeida, J.C., Llerena Junior, J.C. and Reis, D.F. (1992) RAPADILINO syndrome. Am J Med Genet, 44, 716–719.

75. Yakovleva, L. and Shuman, S. (2012) Mycobacterium smegmatis SftH exemplifies a distinctive clade of superfamily II DNA-dependent ATPases with 3’ to 5’ translocase and helicase activities. Nucleic Acids Res, 40, 7465–7475.

76. Kosugi, S., Hasebe, M., Tomita, M. and Yanagawa, H. (2009) Systematic identification of cell cycle-dependent yeast nucleocytoplasmic shuttling proteins by prediction of composite motifs. Proc Natl Acad Sci, 106, 10171–10176.

77. Budd, M.E., Choe, W. and Campbell, J.L. (2000) The nuclease activity of the yeast DNA2 protein, which is related to the RecB-like nucleases, is essential *in vivo*. J. Biol. Chem., 275, 16518–16529.

78. Levikova, M., Klaue, D., Seidel, R. and Cejka, P. (2013) Nuclease activity of Saccharomyces cerevisiae Dna2 inhibits its potent DNA helicase activity. Proc Natl Acad Sci U S A, 110, E1992–2001.

79. Sarek, G., Vannier, J.B., Panier, S., Petrini, J.H.J. and Boulton, S.J. (2015) TRF2 recruits RTEL1 to telomeres in S phase to promote t-loop unwinding. Mol Cell, 57, 622–635.

80. Vannier, J.B., Pavicic-Kaltenbrunner, V., Petalcorin, M.I., Ding, H. and Boulton, S.J. (2012) RTEL1 Dismantles T Loops and Counteracts Telomeric G4-DNA to Maintain Telomere Integrity. Cell, 149, 795–806.

81. Satoh, T., Kato, H., Kumagai, Y., Yoneyama, M., Sato, S., Matsushita, K., Tsujimura, T., Fujita, T., Akira, S. and Takeuchi, O. (2010) LGP2 is a positive regulator of RIG-I- and MDA5-mediated antiviral responses. Proc Natl Acad Sci U S A, 107, 1512–1517.

82. Bruns, A.M., Pollpeter, D., Hadizadeh, N., Myong, S., Marko, J.F. and Horvath, C.M. (2013) ATP hydrolysis enhances RNA recognition and antiviral signal transduction by the innate immune sensor, laboratory of genetics and physiology 2 (LGP2). J Biol Chem, 288, 938–946.

83. Uchikawa, E., Lethier, M., Malet, H., Brunel, J., Gerlier, D. and Cusack, S. (2016) Structural Analysis of dsRNA Binding to Anti-viral Pattern Recognition Receptors LGP2 and MDA5. Mol Cell, 62, 586–602.

84. Liu, Y. (2010) Rothmund-Thomson syndrome helicase, RECQ4: On the crossroad between DNA replication and repair. DNA Repair (Amst), 9, 325–330.

85. Xu, X. and Liu, Y. (2009) Dual DNA unwinding activities of the Rothmund-Thomson syndrome protein, RECQ4. Embo J, 28, 568–577.

86. Fang, H., Nie, L., Chi, Z., Liu, J., Guo, D., Lu, X., Hei, T.K., Balajee, A.S. and Zhao, Y. (2013) RecQL4 helicase amplification is involved in human breast tumorigenesis. PLoS One, 8, e69600.

87. Ho, S.K.L., Tong, G.P.Y., Leung, L.T., Cheng, S.S.W., Fu, E.C.H., Lo, I.F.M., Liu, A.P.Y. and Luk, H.M. (2025) RECQL4-related Rothmund-Thomson syndrome: A case series and literature review. Cancer Genet, **292-293**, 131–136.

88. Li, J., Jin, J., Liao, M., Dang, W., Chen, X., Wu, Y. and Liao, W. (2018) Upregulation of RECQL4 expression predicts poor prognosis in hepatocellular carcinoma. Oncol Lett, 15, 4248–4254.

89. Mo, D., Zhao, Y. and Balajee, A.S. (2018) Human RecQL4 helicase plays multifaceted roles in the genomic stability of normal and cancer cells. Cancer Lett, 413, 1–10.

90. Siitonen, H.A., Sotkasiira, J., Biervliet, M., Benmansour, A., Capri, Y., Cormier-Daire, V., Crandall, B., Hannula-Jouppi, K., Hennekam, R., Herzog, D. et al. (2009) The mutation spectrum in RECQL4 diseases. Eur J Hum Genet, 17, 151–158.

