## Supplementary Materials for "DNA binding contributes to dominant-negative effects of a catalytically inactive RecQ4-family helicase"

**Supplementary files**

**Supplementary File S1.** AlphaFold3 predictions of the *S. cerevisiae* Hrq1 structure containing two Zn^2+^ ions.

**Supplementary File S2.** List of human RECQL4 mutations in the Catalogue Of Somatic Mutations In Cancer (COSMIC) database (1).

**Supplementary tables**

**Table S1. Yeast strains used in this study.**

| **Name** | **Genotype** |
| --- | --- |
| YPH499 | *MATa ura3-52 lys2-801_amber ade2-101_ochre trp1Δ63 his3Δ200 leu2Δ1* |
| MBY520 | *MATa ura3-52 lys2-801_amber ade2-101_ochre trp1Δ63 his3Δ200 leu2Δ1 HRQ1:3xFLAG-His3MX6* |
| MBY745 | *MATa ura3-52 lys2-801_amber ade2-101_ochre trp1Δ63 his3Δ200 leu2Δ1 HRQ1::His3MX6* |
| MBY746 | *MATa ura3-52 lys2-801_amber ade2-101_ochre trp1Δ63 his3Δ200 leu2Δ1 pso2::TRP1* |
| MBY748 | *MATa ura3-52 lys2-801_amber ade2-101_ochre trp1Δ63 his3Δ200 leu2Δ1 HRQ1::hrq1-K318A-His3MX6* |
| MBY945 | *MATa ura3-52 lys2-801_amber ade2-101_ochre trp1Δ63 his3Δ200 leu2Δ1 ylr297w::NatMX* |
| MBY946 | *MATa ura3-52 lys2-801_amber ade2-101_ochre trp1Δ63 his3Δ200 leu2Δ1 tda6::NatMX* |
| MBY947 | *MATa ura3-52 lys2-801_amber ade2-101_ochre trp1Δ63 his3Δ200 leu2Δ1 fmp48::NatMX* |
| MBY959 | *MATa ura3-52 lys2-801_amber ade2-101_ochre trp1Δ63 his3Δ200 leu2Δ1 HRQ1::His3MX6 ylr297w::NatMX* |
| MBY960 | *MATa ura3-52 lys2-801_amber ade2-101_ochre trp1Δ63 his3Δ200 leu2Δ1 HRQ1::His3MX7 tda6::NatMX* |
| MBY961 | *MATa ura3-52 lys2-801_amber ade2-101_ochre trp1Δ63 his3Δ200 leu2Δ1 HRQ1::His3MX8 fmp48::NatMX* |
| MBY1153 | *MATa ura3-52 lys2-801_amber ade2-101_ochre trp1Δ63 his3Δ200 leu2Δ1 chl1::His3MX6* |
| MBY1154 | *MATa ura3-52 lys2-801_amber ade2-101_ochre trp1Δ63 his3Δ200 leu2Δ1 ylr297w::NatMX chl1::His3MX6* |
| MBY1155 | *MATa ura3-52 lys2-801_amber ade2-101_ochre trp1Δ63 his3Δ200 leu2Δ1 tda6::NatMX chl1::His3MX6* |
| MBY1156 | *MATa ura3-52 lys2-801_amber ade2-101_ochre trp1Δ63 his3Δ200 leu2Δ1 fmp48::NatMX chl1::His3MX6* |
| MBY1163 | *MATα ura3-52 lys2-801_amber ade2-101_ochre trp1Δ63 his3Δ200 leu2Δ1 hxt13::URA3 HRQ1::His3MX6::Hrq1DN-His6(NatMX)* |
| MBY1164 | *MATα ura3-52 lys2-801_amber ade2-101_ochre trp1Δ63 his3Δ200 leu2Δ1 hxt13::URA3 HRQ1::His3MX6::Hrq1DN-K318A-His6(NatMX)* |
| MBY1180 | *MATa ura3-52 lys2-801_amber ade2-101_ochre trp1Δ63 his3Δ200 leu2Δ1 HRQ1::hrq1-K318A-3xFLAG-hphMX6* |
| MBY1182 | *MATa ura3-52 lys2-801_amber ade2-101_ochre trp1Δ63 his3Δ200 leu2Δ1 HRQ1::URA3::hrq1-K318AP732S-3xFLAG -hphMX6* |
| MBY1183 | *MATa ura3-52 lys2-801_amber ade2-101_ochre trp1Δ63 his3Δ200 leu2Δ1 HRQ1::URA3::Hrq1-K318AC797Y-3xFLAG -hphMX6* |
| MBY1210 | *MATa ura3-52 lys2-801_amber ade2-101_ochre trp1Δ63 his3Δ200 leu2Δ1 HRQ1::URA3::Hrq1-K318AR739A-3xFLAG -hphMX6* |
| MBY1211 | *MATa ura3-52 lys2-801_amber ade2-101_ochre trp1Δ63 his3Δ200 leu2Δ1 HRQ1::HIS3::Hrq1-R739A-3xFLAG -hphMX6* |
| MBY1212 | *MATa ura3-52 lys2-801_amber ade2-101_ochre trp1Δ63 his3Δ200 leu2Δ1 HRQ1::HIS3::Hrq1-K318AA1006P-3xFLAG -hphMX6* |
| MBY1221 | *MATa ura3-52 lys2-801_amber ade2-101_ochre trp1Δ63 his3Δ200 leu2Δ1 HRQ1::HIS3::Hrq1-C797Y-3xFLAG -hphMX6* |
| MBY1222 | *MATa ura3-52 lys2-801_amber ade2-101_ochre trp1Δ63 his3Δ200 leu2Δ1 HRQ1::HIS3::Hrq1-A1006P-3xFLAG -hphMX6* |
| MBY1228 | *MATa ura3-52 lys2-801_amber ade2-101_ochre trp1Δ63 his3Δ200 leu2Δ1 HRQ1::HIS3::Hrq1-P732S-3xFLAG -hphMX6* |

**Table S2. Yeast expression plasmids used in this study.**

| **Name** | **Gene involved** | **Markers** | **Notes** |
| --- | --- | --- | --- |
| pMB346 | Empty vector | *Amp^R^, LEU2* | Contains the regions up- and downstream of the *HRQ1* gene for transcriptional control; *CEN* vector |
| pMB389 | *hrq1-K318A* | *Amp^R^, LEU2* | *3xFLAG-hrq1-K318A* cloned into pMB346 |
| pMB390 | *HRQ1* | *Amp^R^, LEU2* | *3xFLAG-HRQ1* cloned into pMB346 |
| pMB911 | *hrq1-K318A-4CS* | *Amp^R^, LEU2* | *3xFLAG-hrq1-K318A-4CS* cloned into pMB346 |
| pMB912 | *hrq1-K318A-C981Y* | *Amp^R^, LEU2* | *3xFLAG-hrq1-K318A-C981Y* cloned into pMB346 |
| pMB913 | *hrq1-K318A-C985Y* | *Amp^R^, LEU2* | *3xFLAG-hrq1-K318A-C985Y* cloned into pMB346 |
| pMB914 | *hrq1-K318A-C988Y* | *Amp^R^, LEU2* | *3xFLAG-hrq1-K318A-C988Y* cloned into pMB346 |

**Table S3.**

| **Mutation** | **ESM-1v score** | **ESM-1v Class^1^** | **FoldX ΔΔG** | **FoldX Class^2^** | **MAESTRO ΔΔG** | **MAESTRO Class^3^** |
| --- | --- | --- | --- | --- | --- | --- |
| P48A | -3.293 | Likely Benign | 1.59335 | Destabilizing | 0.258328839 | Neutral/Marginal |
| P48T | -3.109 | Likely Benign | 2.0385 | Destabilizing | 0.301346306 | Neutral/Marginal |
| Q121K | -2.752 | Likely Benign | -0.064129 | Neutral | -0.351655248 | Neutral/Marginal |
| N122Y | -4.188 | Likely Benign | -0.756114 | Neutral | -0.057515598 | Neutral/Marginal |
| F181L | -2.453 | Likely Benign | 0.420712 | Neutral | 0.175285984 | Neutral/Marginal |
| K318A | -14.939 | Highly Deleterious | ND | ND | ND | ND |
| N409K | -1.729 | Likely Benign | -0.1675 | Neutral | -0.223560219 | Neutral/Marginal |
| R442C | -11.530 | Highly Deleterious | 5.57742 | Destabilizing | 0.195869629 | Neutral/Marginal |
| R442G | -12.554 | Highly Deleterious | 6.6216 | Destabilizing | 0.49687732 | Neutral/Marginal |
| H448N | -2.927 | Likely Benign | 0.546095 | Neutral | 0.035012652 | Neutral/Marginal |
| L545F | -5.726 | Likely Benign | 3.82427 | Destabilizing | 0.193413738 | Neutral/Marginal |
| V641G | -10.671 | Highly Deleterious | 5.00139 | Destabilizing | 0.373129807 | Neutral/Marginal |
| H683D | -10.608 | Highly Deleterious | 2.83449 | Destabilizing | 0.130731288 | Neutral/Marginal |
| L691P | -9.065 | Highly Deleterious | 6.41477 | Destabilizing | 0.415424782 | Neutral/Marginal |
| P732S | -4.292 | Likely Benign | 4.85077 | Destabilizing | 0.147143382 | Neutral/Marginal |
| R739A | -9.827 | Likely Deleterious | ND | ND | ND | ND |
| R739E | -10.867 | Highly Deleterious | ND | ND | ND | ND |
| A747P | -6.854 | Likely Deleterious | 0.402599 | Neutral | 0.095300214 | Neutral/Marginal |
| Q780K | -3.728 | Likely Benign | 1.06484 | Destabilizing | 0.288459096 | Neutral/Marginal |
| A900E | -7.674 | Likely Deleterious | 8.84606 | Destabilizing | 0.188121894 | Neutral/Marginal |
| S972N | -6.929 | Likely Deleterious | 5.13695 | Destabilizing | 0.177706969 | Neutral/Marginal |
| C979F | -6.567 | Likely Deleterious | 8.3215 | Destabilizing | 0.127491906 | Neutral/Marginal |
| C979L | -6.408 | Likely Deleterious | -1.7857 | Neutral | 0.208867742 | Neutral/Marginal |
| C979W | -8.211 | Likely Deleterious | 9.71848 | Destabilizing | 0.123927733 | Neutral/Marginal |
| C979Y | -6.751 | Likely Deleterious | 8.30851 | Destabilizing | 0.037436121 | Neutral/Marginal |
| G1005C | -7.568 | Likely Deleterious | 6.29963 | Destabilizing | -0.00195762 | Neutral/Marginal |
| A1006P | -9.207 | Likely Deleterious | 5.12607 | Destabilizing | 0.272077409 | Neutral/Marginal |

**^1^** The ESM-1v scores were classified as follows: > -6, Likely Benign; -6 to -10, Likely Deleterious; and < -10, Highly Deleterious.

^2^ The FoldX ΔΔG values were classified as follows: < 1.0 kcal/mol, neutral; and ≥ 1.0 kcal/mol, destabilizing.

^3^ The MAESTRO ΔΔG values were classified as follows: < -0.5, Stabilizing; -0.5-0.5, Neutral/Marginal; 0.5-1.0, Mildly Destabilizing; 1.0-2.0, Moderately Destabilizing; and > 2.0, Highly Destabilizing.

**Supplementary Figure Legends**

**Figure S1. The FMP48, TDA6, and YLR297W genes do not participate in DEB ICL repair in *S. cerevisiae*.** Cells of the indicated genotypes were grown in liquid culture, diluted to OD_600_ = 1, and 10-fold serial dilutions were spotted onto plates containing rich medium (YPD) or YPD supplemented with the indicated concentration of DEB. The plates were then incubated for 2 (YPD and YPD + 50 mg/mL DEB) or 3 (YPD + 100 mg/mL DEB and YPD + 150 mg/mL DEB) days in the dark prior to imaging.

**Figure S2. Second-site suppressors of *hrq1-K318A* DEB sensitivity.** Cells of the indicated genotypes were grown overnight, diluted to OD_600_ = 1, ten-fold serially diluted, and then 5 μL of each dilution was spotted onto YPD or YOD supplemented with DEB. The plates were incubated at 30°C prior to imaging with Bio-Rad ChemiDoc MP system.

**Figure S3. Western blot analysis of Hrq1 and mutant Hrq1 protein levels.** This is the un-truncated blot, parts of which are shown in Figures 2C and 5C in the main text. The amounts of mutant proteins were compared to WT using one-way ANOVA. NS, *p* > 0.01; *, *p* < 0.01.

**Figure S4. The *hrq1-K318A* allele is a dominant negative.** A) Wild-type Hrq1 expression can rescue the diepoxybutane (DEB) sensitivity of the *hrq1Δ* strain (MBY745). Cells lacking the *HRQ1* gene (*hrq1Δ*) were transformed with either an empty single-copy *LEU2*-marked vector (Vector) or the same vector encoding *HRQ1* under the transcriptional control of its native promoter. Overnight cultures of these strains were diluted to OD_600_ = 1, serially diluted 10-fold, and 10 μL of each dilution was plated on leucine drop-out medium (-Leu) and -Leu supplemented with DEB. The plates were incubated at 30°C for 2-5 days prior to imaging with a flatbed scanner. B) Hrq1 expression fails to rescue the DEB sensitivity of the *hrq1-K318A* mutant (MBY748). The experiment was performed as in (A), but the parental strain carried the *hrq1-K318A* allele at its native locus.

**Figure S5. CLUSTAL alignment of the Hrq1, RECQL4, and MrfA sequences.** The primary sequences of *S. cerevisiae* Hrq1 (ScHrq1; WNF20109.1), *Homo sapiens* RECQL4 (HsRECQL4; BAA86899.1), and *Bacillus subtilis* MrfA (BsMrfA; WP_015383941.1) were aligned using the CLUSTAL Omega (v1.2.4) algorithm, which utilizes a seeded guide tree and HMM profile-profile techniques to generate accurate alignments of divergent sequences. Second-site suppressor mutations in Hrq1 are noted in bold red font, and if the residue is conserved in RECQL4 and/or MrfA, the corresponding sites are also in bold red font. Asterisks (*) denote positions with a fully conserved residue. Colons (:) indicate conservation between groups of strongly similar properties (roughly > 0.5 in the Gonnet PAM 250 matrix). Periods (.) indicate conservation between groups of weakly similar properties (roughly ≤ 0.5 in the Gonnet PAM 250 matrix). Dashes (-) represent gaps introduced to maximize the alignment of conserved motifs.

**Figure S6.** **Second-site suppressors map to all Hrq1 domains.** A) The three-dimensional structure of Hrq1 was predicted using the AlphaFold Server (<https://alphafoldserver.com>) and included two Zn^2+^ ions (ipTM = 0.89 and pTM = 0.68). The structure is coloured according to the per-atom confidence estimate (plDDT), where a higher value indicates higher confidence. B) Predicted alignment error (PAE) plot showing the approximate positions of the Cdt1-like, helicase, RQC, and RHCD domains. C) Predicted Hrq1 structure with the suppressor mutations mapped. The four domains of the enzyme are marked on the left. Red residues denote substitution mutations, and blue denotes the sites of frameshifts/premature stop codons. The C979 and A1006 sites are labelled in the central panel.

**Figure S7. A1006 is predicted to lie in close proximity to the 4-Cys motif.** In the AlphaFold 3 model of the Hrq1 structure, residue A1006 is ~8 Å away from the closest Cys (C988) in the putative Zn^2+^-binding motif.

**Figure S8. Denaturing Hrq1 destroys its oligomeric forms.** Mass photometry traces of the in-solution molecular mass of Hrq1-K318A,R739A (top) and the same protein denatured with 1% SDS and a 10-min incubation at 95°C (bottom). The native protein exists as monomers, dimers, trimers, and tetramers in solution, but the denatured protein only displays a peak consistent with the size of a monomer.

**Figure S9. Truncating the N-terminus of Hrq1 suppresses the ICL sensitivity of the K318A mutant.** Strains of the indicated genotypes were grown overnight, diluted to OD600 = 1, serially diluted tenfold, and then 5 μL of each dilution was spotted on to YPD or YPD supplemented with DEB.

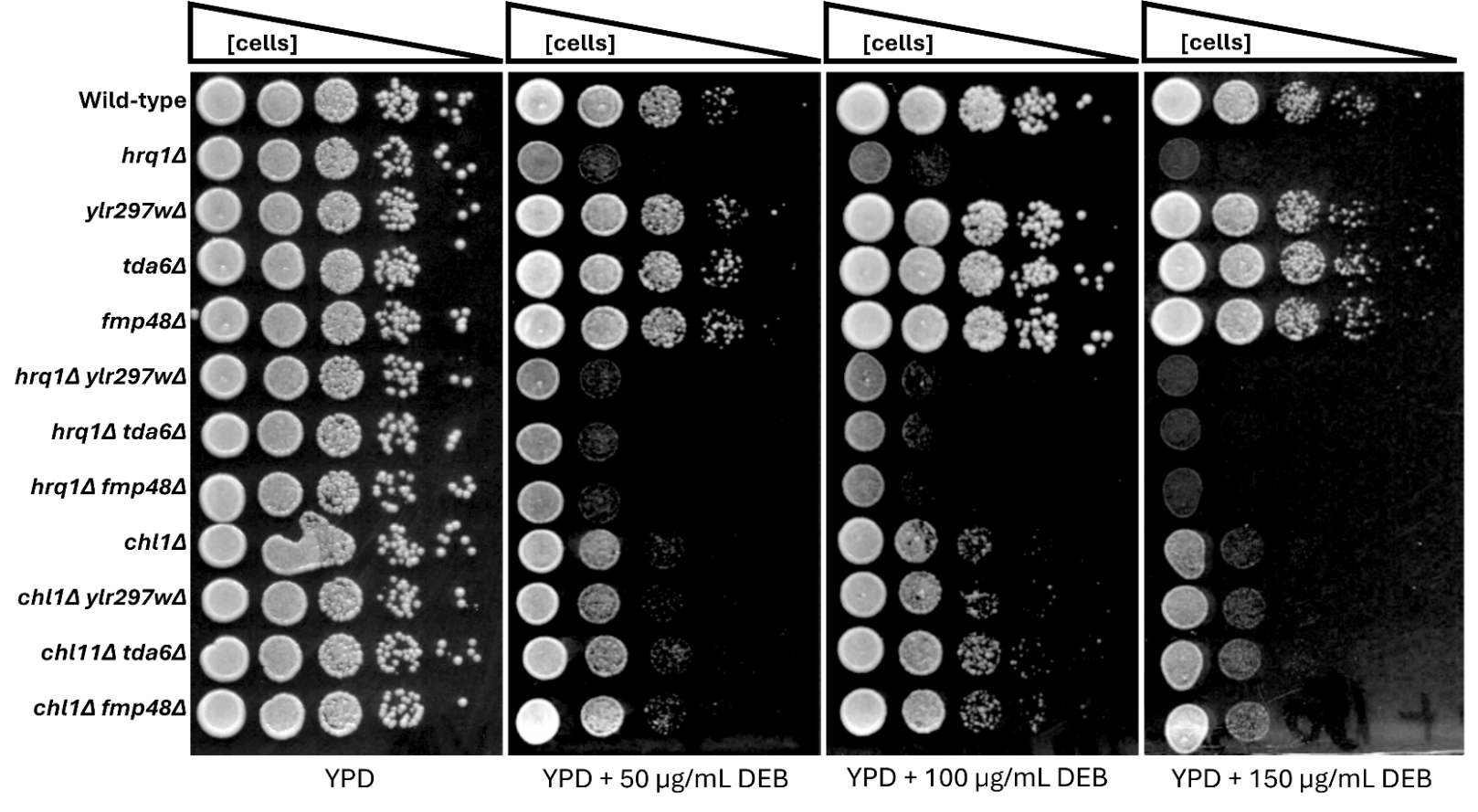

**Figure S1.**

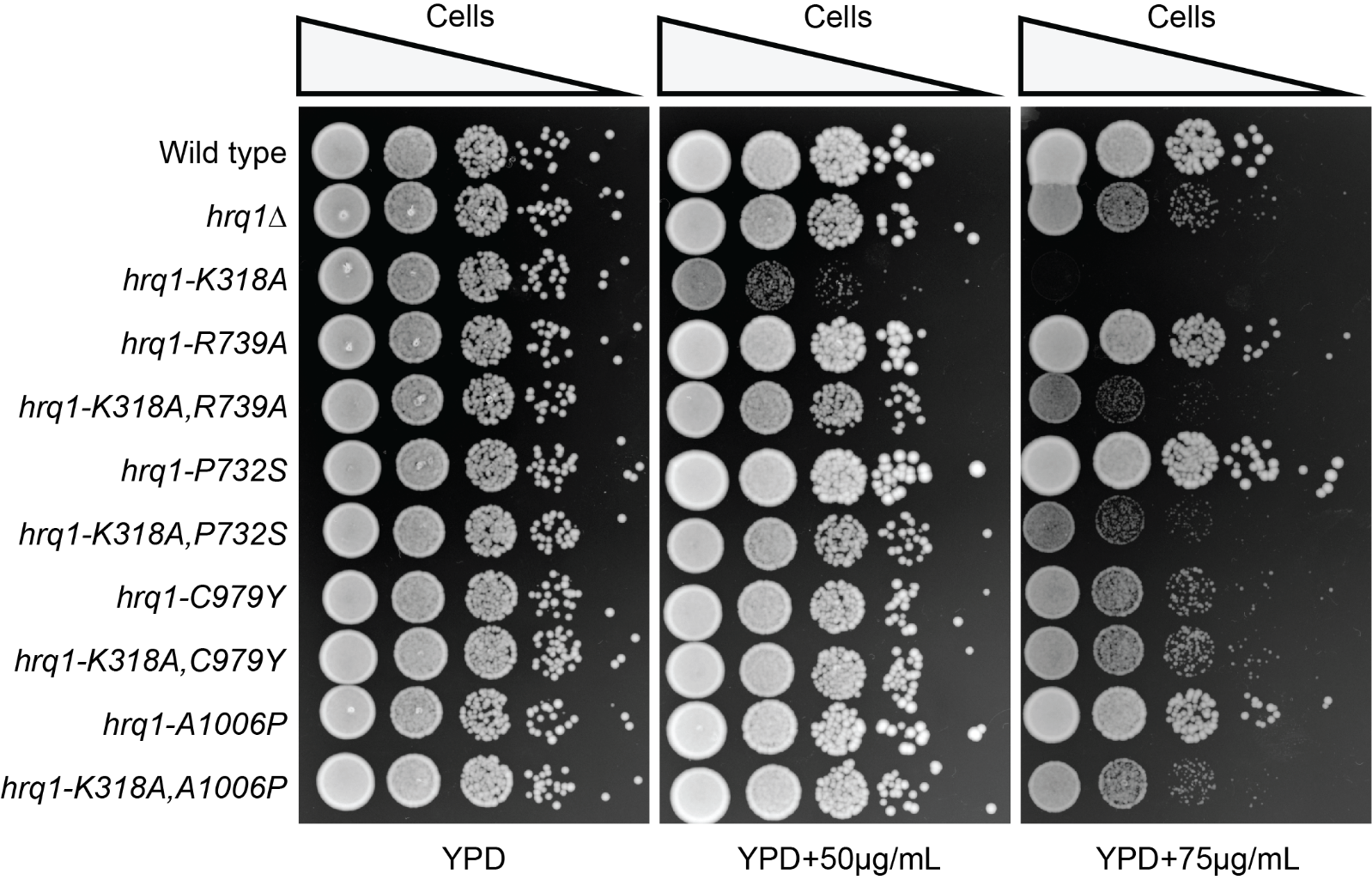

**Figure S2.**

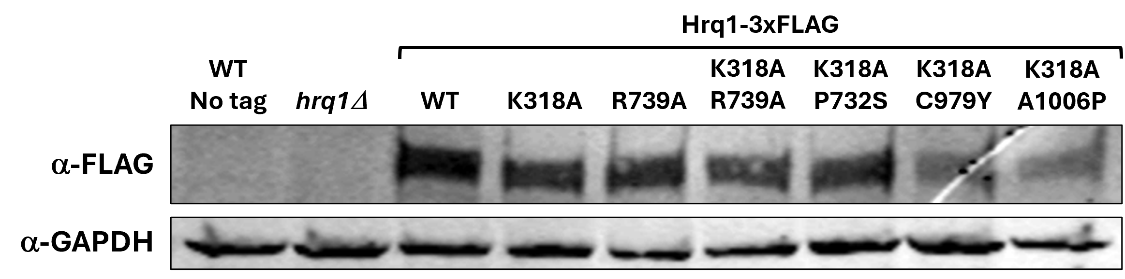

**Figure S3**

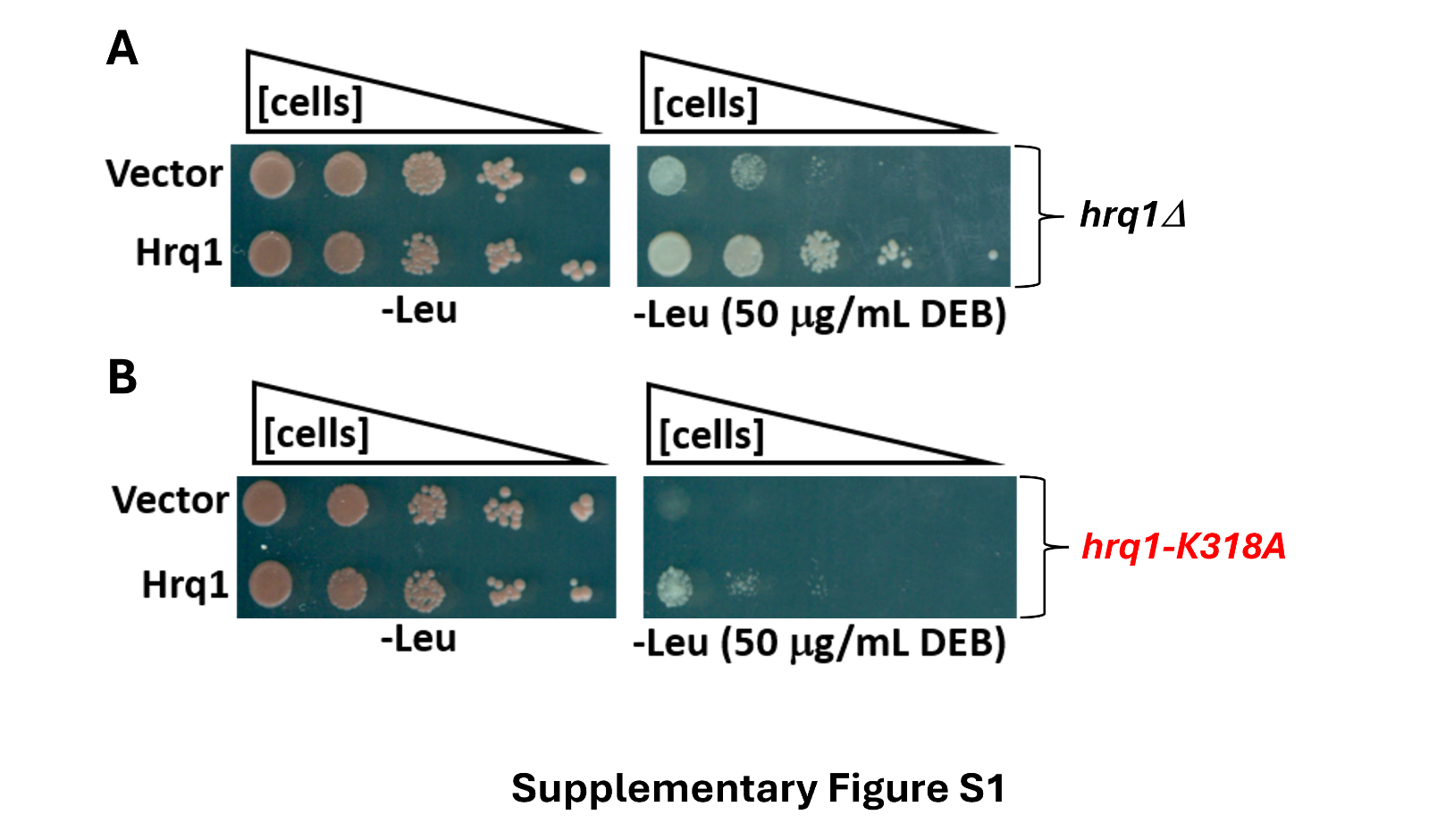

**Figure S4.**

```
ScHrq1        ------------------------------------------------------------
HsRECQL4      MERLRDVRERLQAWERAFRRQRGRRPSQDDVEAAPEETRALYREYRTLKRTTGQAGGGLR
BsMrfA        ------------------------------------------------------------


ScHrq1        --------------------------------------------------------MEEG
HsRECQL4      SSESLPAAAEEAPEPRCWGPHLNRAATKSPQPTPGRSRQGSVPDYGQRLKANLKGTLQAG
BsMrfA        ------------------------------------------------------------


ScHrq1        PIKKK------------LKSAGQGSGKTDAFR---------------------NFE-QFF
HsRECQL4      PALGRRPWPLGRASSKASTPKPPGTGPVPSFAEKVSDEPPQLPEPQPRPGRLQHLQASLS
BsMrfA        ------------------------------------------------------------

                                 P48A/T
ScHrq1        FRLNTLYT-F-LICRKHVVPTFKT---LCGPIETALKRTVTKEDLAMVMALMPRECVFKY
HsRECQL4      QRLGSLDPGWLQRCHS-EVPDFLGAPKACRPDL-------GSEES---QLLIPGES----
BsMrfA        ------------------------------------------------------------

                                                  N122Y
ScHrq1        IDENQIYTETKIFDFNNGGFQQKENDIFELKDVDDQNQTQKSTQLLIFEFIDGTMQRSWS
HsRECQL4      ----------AVLGPGA-GSQGPEASAFQEVSIRVGSP---------QPSSSGGEKRRWN
BsMrfA        ------------------------------------------------------------


ScHrq1        ASDRFSQI-----------------------------------------KIPTYTTEEMK
HsRECQL4      EEPWESPAQVQQESSQAGPPSEGAGAVAVEEDPPGEPVQAQPPQPCSSPSNPRYHGLSPS
BsMrfA        ------------------------------------------------------------

                                F181L
ScHrq1        K--MISKREALFKSRLREFILEKEKANLDPFSELTNLAQKYIPRERDYEDPIEA--MMKA
HsRECQL4      SQARAGKAEGT--APLHIFP-RLARHDRGNYVRLNMKQKHYV-RGRALRSRLLRKQAWKQ
BsMrfA        ---------------------------------------------------------MKK
                                                                        *

ScHrq1        KQESNEMSIPNYSNNSVITTIPQMIEKLKSTEFYASQIKHCFTIPSRTAKY---------
HsRECQL4      KWRKKG---ECFGGG-------GATVTTKESCFLNEQFDHWAAQCPRPASEEDTDAVGPE
BsMrfA        K------------------SLTELISDLKGNEN----VVNWHEIEPREAKT---------
              *                           * .      . :      * *.

ScHrq1        ---------------------------KGLCFEL---APEVYQGMEHENFYSHQADAINS
HsRECQL4      PLVPSPQPVPEVPSLDPTVLPLYSLGPSGQLAETPAEVFQALEQLGHQAFRPGQERAVMR
BsMrfA        ---------------------------RPMPESIDERIKAALSKRGIDELYTHQYSAFQY
                                              .       . .    : :   *  *.

ScHrq1        LHQGENVIITTSTSSGKSLIYQLAAIDLLLKDPESTFMYIFPTKALAQDQKRAFKVILSK
HsRECQL4      ILSGISTLLVLPTGAGKSLCYQLPALLYSRRSPCLT-LVVSPLLSLMDDQVSGLPPCLKA
BsMrfA        VQKGESIVTVTPTASGKTLCYNLPVLQSIAQDETNRALYLFPTKALAQDQKSELNEIIDE
              : .* . : .  *.:**:* *:* .:    :.     : : *  :* :**   :   :.
                                                               N409K
ScHrq1        IPELKNAVVDTYDGDTEPEERAYIRK-NARVIFTNPDM-IHTSILPNHANWRHFLYHLKL
HsRECQL4      ACIHSGMT-----RKQRESVLQKIRAAQVHVLMLTPEALVGAGGLPPAAQ----LPPVAF
BsMrfA        MGI--DIKSFTYDGDTSPAIRQKVRK-AGHIVITNPDM-LHSAILPHHTKWVSLFENLKY
                   .        .        :*    :::: .*:  : :. **  ::    :  :
                                    R422C/G
ScHrq1        VVVDELHIYKGLFGSHVALVMRRLLRLCHCFYENSGLQF-ISCSATLKSPVQHMKDMFGI
HsRECQL4      ACIDEAHCLSQWS----HNFRPCYLRVCKVLRERMGVHCFLGLTATATR-----------
BsMrfA        IVIDELHTYRGVFGSHVANVIRRLKRICRFYGS--DPVF-ICTSATIANPKEL-GEQLTG
                :** *            .     *:*:   .  .    :  :**

ScHrq1        NEVTLIHEDGSPTGAKHLVVWN------PPILPQHERKRENFIRESAKILVQL-----IL
HsRECQL4      --------RTASDVAQHLAVAEEPDLHGPAPVPTNLHLSVSMDRDTDQALLTLLQGKRFQ
BsMrfA        KPMRLVDDNGAPSGRKHFVFYN------PPIVNKPLNIRRSATAEVNELAKEF-----LK
                        :    :*:.. :      *  :    .   .   :  :    :     :
                               L545F
ScHrq1        NNVRTIAFCYVRRVCELLMKEVRNI-----FIETGREDLVTEVMSYRGGYSASDRRKIER
HsRECQL4      NLDSIIIYCNRREDTERIAALLRTCLHAAWVPGSGGRAPKTTAEAYHAGMCSRERRRVQR
BsMrfA        NKVQTIVFARSRVRVEIILSHIQEL-----VK---KEIGTKSIRGYRGGYLPKERREIER
              *    * :.  *   * :   ::       .     .   .   .*:.*    :**.::*
                                                                        V641G
ScHrq1        EMFHGNLKAVISTNALELGIDIGGLDAVLMCGFPLSMANFHQQSGRAGRRNNDSLTLVVA
HsRECQL4      AFMQGQLRVVVATVAFGMGLDRPDVRAVLHLGLPPSFESYVQAVGRAGRDGQPAHCHLFL
BsMrfA        GLREGDILGVVSTNALELGVDIGQLQVCVMTGYPGSVASAWQQAGRAGRRHGESLIIMVA
               : .*::  *::* *: :*:*   : . :  * * *. .  *  *****    :   :.
                                                           H683D   L691P
ScHrq1        SDSPVD-----QHYVAHPESLLEVNNFESYQDLVLDFNNILILEGHIQCAAFELPINFER
HsRECQL4      QPQGEDLRELRRHVHADSTDFLAVKRLV------------QRVFPACTCTCTRPPSEQEG
BsMrfA        NSTPID-----QYIVRHPEYFFNRSP----ESARINPENLIILVDHLKCAAYELPFRADE
              .    *     ::   .   ::  .                 :     *:. . * . :

ScHrq1        DKQY-----------------FTESHLRKICV--ERL----------------------H
HsRECQL4      AVGGERPVPKYPPQEAEQLSHQAAPGPRRVCMGHERALPIQLTVQALDMPEEAIETLLCY
BsMrfA        EFGA-----------------MEVSDILEYLQEEAVL----------------------H
                                          .                              :
                             P732S
ScHrq1        HNQDGYHASNRFLPWPSKCV-SLRGGEEDQFAVVDITNGRNI-IIE----EIE-----AS
HsRECQL4      LELHPHHWLELLATTYTHCRLNCPGGPAQLQALAHRCPPLAVCLAQQLPEDPGQGSSSVE
BsMrfA        RNGERYHWASE--SFPASNI-SLRSASQENVVIVDQSDIANVRIIG----EMD-----RF
               : . :*  .      :    .  ..  :  .:..      : :      :
                            Q780K
ScHrq1        RTSFTLYDGGIFIHQGYPYLVKEFNPDERYAKVQRVDVDWVTN-QRDFTDVDPQEIELIR
HsRECQL4      FDMVKLVDSMGWELASVRRALCQLQWDHEPRTGVRRGTGVLVEFSELAFHLRS-PGDLTA
BsMrfA        SAMTLLHDEAIYLHEGVQYQVEKLDWDHKKAYVRKVDVEYYTD-ANLAVQLKVLETD--K
                   * *   :   .    : ::: *..     : ..   .:  .   .:     :

ScHrq1        SLRNSDVPVYFGKIKTTII--------VFGFFKVDKYKRIIDAIETHNPPVIINSKGLWI
HsRECQL4      EEKDQICDFLYGRVQARERQALARLRRTFQAFHSVAFPS----------------CGPCL
BsMrfA        TKEKSRTSLHYGDVTVNAL------PTIFKKIKMTTFEN-IGSGPIHLPEEELHTSAAWL
                ...   . :* : .            *  ::   :                   .  :
                                          A900E
ScHrq1        DMPKYALEICQKKQL-----NVAGAIHGAQHAIMGMLPR---------FIVAGVDEIQTE
HsRECQL4      EQQDEERSTRLKDLLGRYFEEEEGQEPGGMEDAQGPEPGQARLQDWEDQVRCDIR--QFL
BsMrfA        EIKTADEDIGEKTL--------EQLLLGISNVLQHIVPV---------YIMCDRNDVHVV
              :      .   *               *  .      *           : ..    :
                                                               S972N  C979F/L/W/Y
ScHrq1        CKAPEKEFAERQTKRKRPARLIFYDSKGGKYGSGLCVKAFEHIDDIIESSLRRIEECPCS
HsRECQL4      SLRPEEKFSSRAVA-----RIF------------------HG------------------
BsMrfA        SQ--------IKAAHTGLPTIFLYDHYPG--GIGLAEEVFKRFSDINEAAKQLIAHCPCH
              .           .       ::                  .

      G1005C, A1006P
ScHrq1        DGCPDCVAASFCKENSLVLSKPGAQVVLHCILGHSEDSFIDLIKDGPEPNMPEIKVETVI
HsRECQL4      IG-SPCYPAQVYGQDRRFWRK-YLHLSFHALVGLATEELLQVAR----------------
BsMrfA        DGCPSCIGTEIEGIKAKE-------RILQL-LDQMS------------------------
               *   *  :..   .            ::  :.

ScHrq1        PVSEHVNFSDDFKIIDVRRATKDDTHTNEIIKKEI
HsRECQL4      -----------------------------------
BsMrfA        -----------------------------------
```

**Figure S5.**

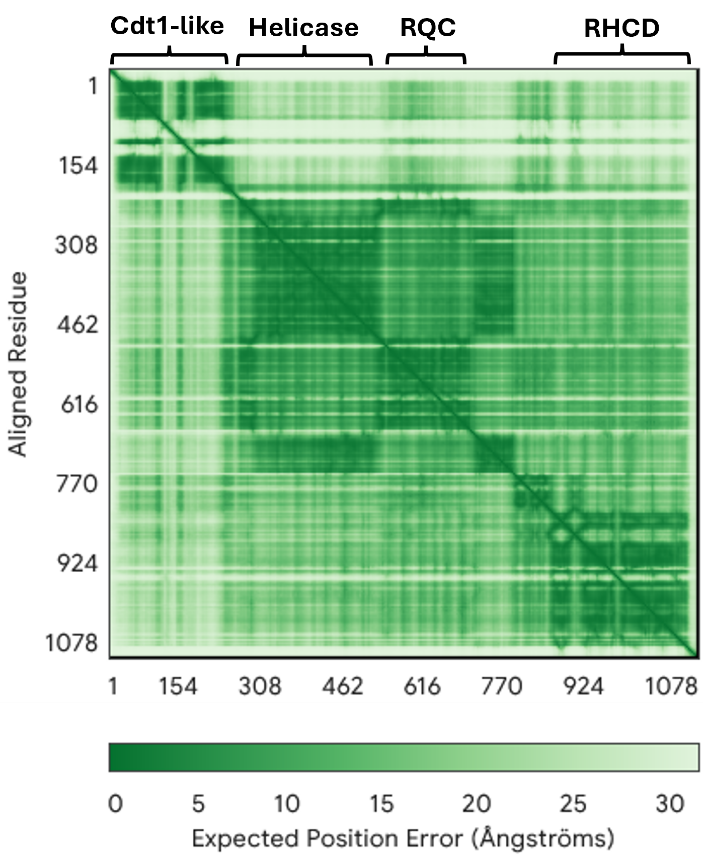

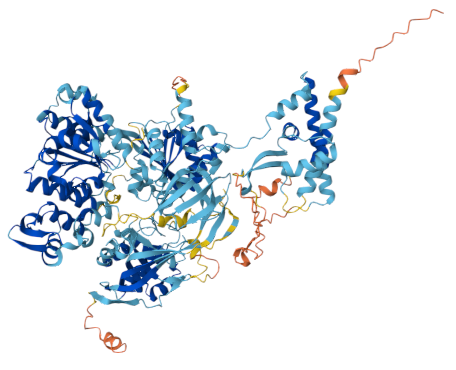
**A	B**

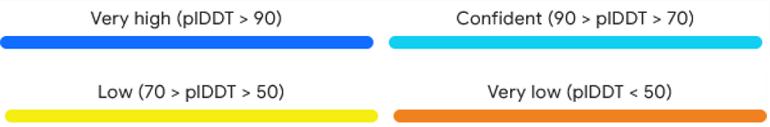

**C**
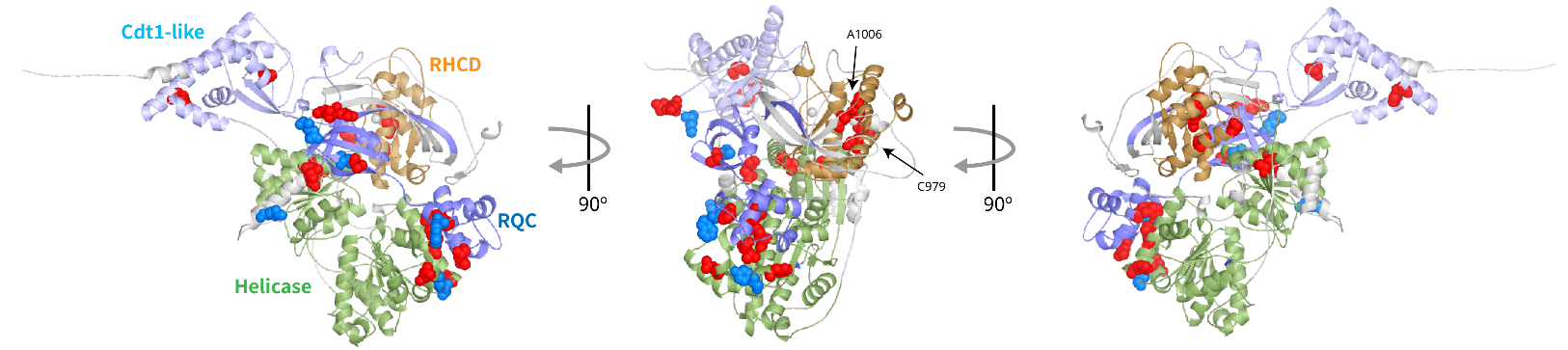

**Figure S6**

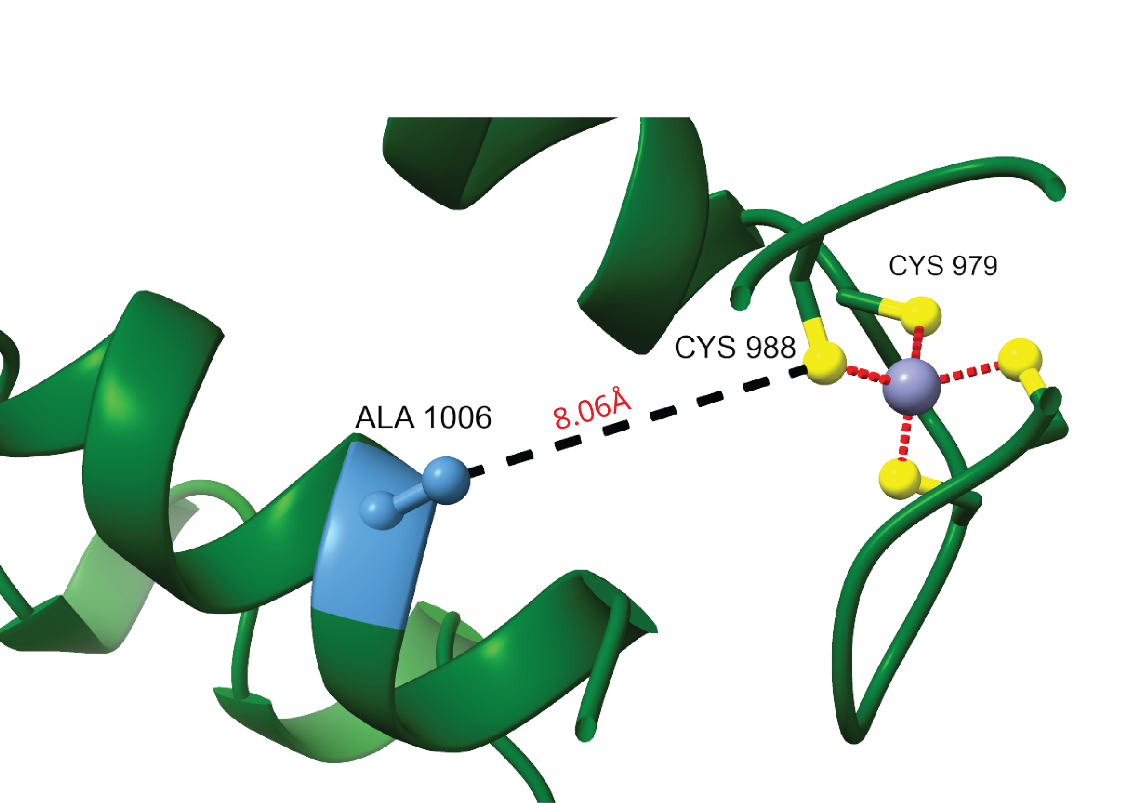

**Figure S7.**

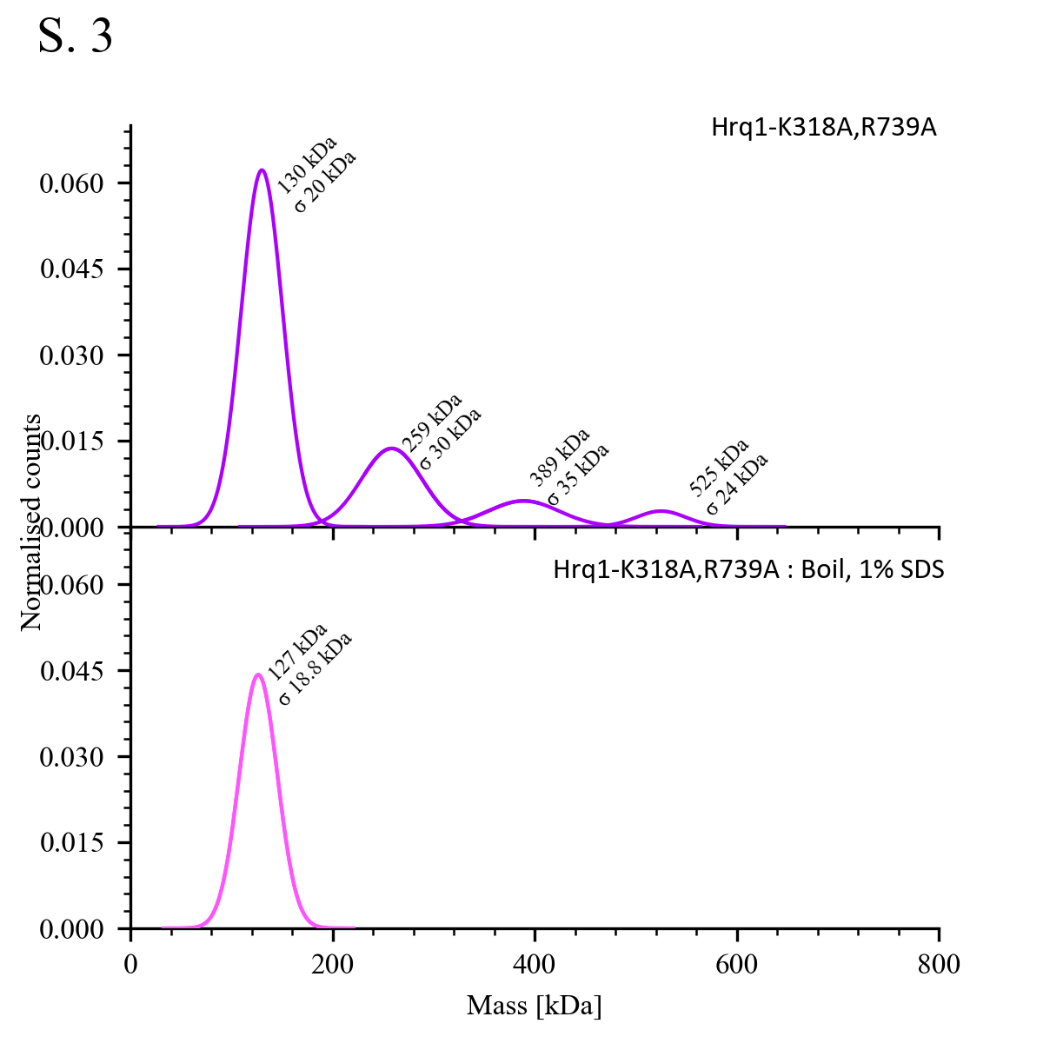

**Figure S8.**

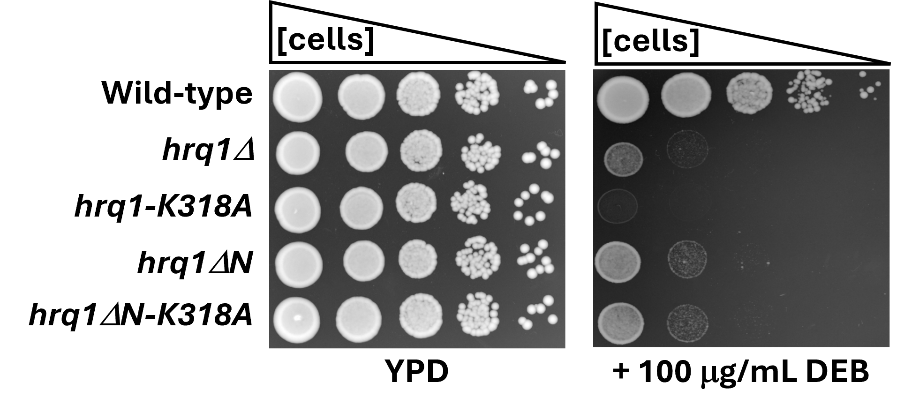

**Figure S9.**

**Supplementary References**

1.	Tate, J.G., Bamford, S., Jubb, H.C., Sondka, Z., Beare, D.M., Bindal, N., Boutselakis, H., Cole, C.G., Creatore, C., Dawson, E. *et al.* (2019) COSMIC: the Catalogue Of Somatic Mutations In Cancer. *Nucleic Acids Res*, **47**, D941-D947.
